# Casein Kinase 1G2 Suppresses Necroptosis-Promoted Testis Aging by Inhibiting Receptor-Interacting Kinase 3

**DOI:** 10.1101/2020.08.17.254318

**Authors:** Dianrong Li, Youwei Ai, Jia Guo, Baijun Dong, Lin Li, Gaihong Cai, She Chen, Dan Xu, Fengchao Wang, Xiaodong Wang

## Abstract

Casein kinases are a large family of intracellular serine/threonine kinases that control a variety of cellular signaling functions. Here we report that a member of casein kinase 1 family, casein kinase 1G2, CSNK1G2, binds and inhibits the activation of receptor-interacting kinase 3, RIP3, thereby attenuating RIP3-mediated necroptosis. The binding of CSNK1G2 to RIP3 is triggered by auto-phosphorylation at serine 211/threonine 215 sites in its C-terminal domain. CSNK1G2-knockout mice showed significantly enhanced necroptosis response and pre-maturing aging of their testis, a phenotype that was rescued by either double knockout of the RIP3 gene or feeding the animal with a RIP1 kinase inhibitor-containing diet. Moreover, CSNK1G2 is also co-expressed with RIP3 in human testis, and the necroptosis activation marker phospho-MLKL was observed in the testis of old (>80) but not young men, indicating that the testis-aging program carried out by the RIP3-mediated and CSNK1G2-attenuated necroptosis is evolutionarily conserved between mice and men.

## INTRODUCTION

RIP3 is an intracellular serine/threonine kinase with key roles in necroptosis, a regulated form of necrotic cell death, and activation of inflammasome for the release of inflammatory cytokines (Christofferson and Yuan, 2010; Vandenabeele et al., 2010; Wallach et al., 2016). During necroptosis, in response of TNF-family of cytokines, Toll-like receptors, and Z-RNAs, RIP3 is activated either by a related kinase RIP1, or adaptor proteins TRIF, or ZBP1/DAI, respectively (Cho et al., 2009; Degterev et al., 2008; He et al., 2009; Zhang et al., 2009; Zhang et al., 2020). Activated RIP3 phosphorylates the mixed lineage kinase-domain like protein, MLKL, to trigger its oligomerization and translocation from the cytosol to membranes including plasma membrane for their disruption (Cai et al., 2014; Chen et al., 2014; Sun et al., 2012; Wang et al., 2014). Necroptosis is actively suppressed by caspase-8-mediated cleavage of RIP1 and RIP3, whose upstream activation pathway is often shared between caspase-8 and RIP1 kinase. Embryonic lethality in caspase-8 knockout mice is rescued by double knockout RIP3 or MLKL; and cellular necroptosis induction by TNF-α or TLRs are dramatically enhanced if caspase-8 activity is suppressed (Dannappel et al., 2014; Dillon et al., 2014; Gunther et al., 2011; He et al., 2009; Kaiser et al., 2011; Newton et al., 2019; Oberst et al., 2011; Rickard et al., 2014; Takahashi et al., 2014). RIP3 was also reported to be negatively regulated by the phosphatase Ppm1b but the effect seems minor and the *in vivo* validation has yet to be obtained (Chen et al., 2015).

One of the important physiological function of necroptosis is to promote the aging of testis in mice (Li et al., 2017). The necroptosis activation marker, the phosphorylated MLKL, has been observed in spermtagonium stem cells and Sertoli cells in the seminiferous tubules of old (>18 months) but not young mouse testis. Knockout RIP3, or MLKL, or feeding mice with a RIP1 kinase inhibitor-containing diet, blocks necroptosis from occurring in mouse testis, and allows mice to maintain the youthful morphological features and the function of male reproductive system to advanced ages, when age-matched wild type mice had lost their reproductive function (Li et al., 2017).

Casein kinases are a large family of intracellular serine/threonine kinases that control a variety of cellular signaling functions that include the circadian clock, Wnt receptor activation, μ opioid receptor modulation, DNA repair, and hypoxia response (Davidson et al., 2005; Elyada et al., 2011; Etchegaray et al., 2009; Goldberg et al., 2017; Pangou et al., 2016). We found one of the casein kinase 1 family members, casein kinase 1G2, CSNK1G2, binds to RIP3 and the binding inhibits RIP3 kinase activity. Interestingly, CSNK1G2 expresses at the highest level in mouse testis, and whose expression pattern overlaps with that of RIP3. Knocking out CSNK1G2, in mouse or multiple cell lines including cell lines derived from spermatocyte and Sertoli cells significantly enhanced necroptosis response and the CSNK1G2 knockout mice showed pre-maturing aging of their testis. Our results demonstrate that CSNK1G2 is a major negative regulator of necroptosis via prevention of RIP3 activation.

## RESULTS

### CSNK1G2 Negatively Regulates Necroptosis by Binding to RIP3

In a course of investigating RIP3-interacting proteins, we found that several members of the casein kinase 1 family were among the proteins co-precipitated with RIP3 kinase (Figure S1A). The effect of CSNK1 members on RIP3 kinase activity was then assessed by co-expressing each member with RIP3 in human embryo kidney 293T cells, and the RIP3 kinase activity was measured by probing the serine 227 auto-phosphorylation of RIP3, an event critical for RIP3 to recruit its substrate MLKL (Li et al., 2015; Sun et al., 2012). Among the casein kinase family members, CSNK1D1, CSNK1G2, and CSNK1E suppressed serine 227 phosphorylation on RIP3 (Figure S1B). In particular, CSNK1G2, but not its closest family members CSNK1G1 and CSNK1G3, showed the most potent suppression of RIP3 kinase activity (Figure S1C). Two kinase-dead mutants, CSNK1G2(K75A) and CSNK1G2(D165N), failed to suppress RIP3 kinase activity (Figure 1A), indicating that the kinase activity of CSNK1G2 is required for its function in suppressing RIP3. Consistently, knockout CSNK1G2 in mouse embryonic fibroblasts, MEFs, significantly accelerated MEF necroptosis induced by the combination of <u>T</u>NF-α (T), a <u>S</u>mac mimetic (S), and a pan-caspase inhibitor <u>Z</u>-VAD-fmk (Z) (Figure 1B). The enhanced necroptosis was mitigated by re-introducing wild type CSNK1G2 into the CSNK1G2 knockout MEFs, but a similar level of K75A kinase-dead mutant did not restore the necroptosis inhibition activity (Figure 1B). In addition to TSZ, MEFs with their CSNK1G2 knocked out also showed enhanced cell death when treated with death-inducing cytokine TRAIL plus a Smac mimetic and z-VAD (TRAIL/S/Z), or lipopolysaccharide (LPS) plus a Smac mimetic and z-VAD (LPS/S/Z) (Figure S2A).

**Figure 1.**
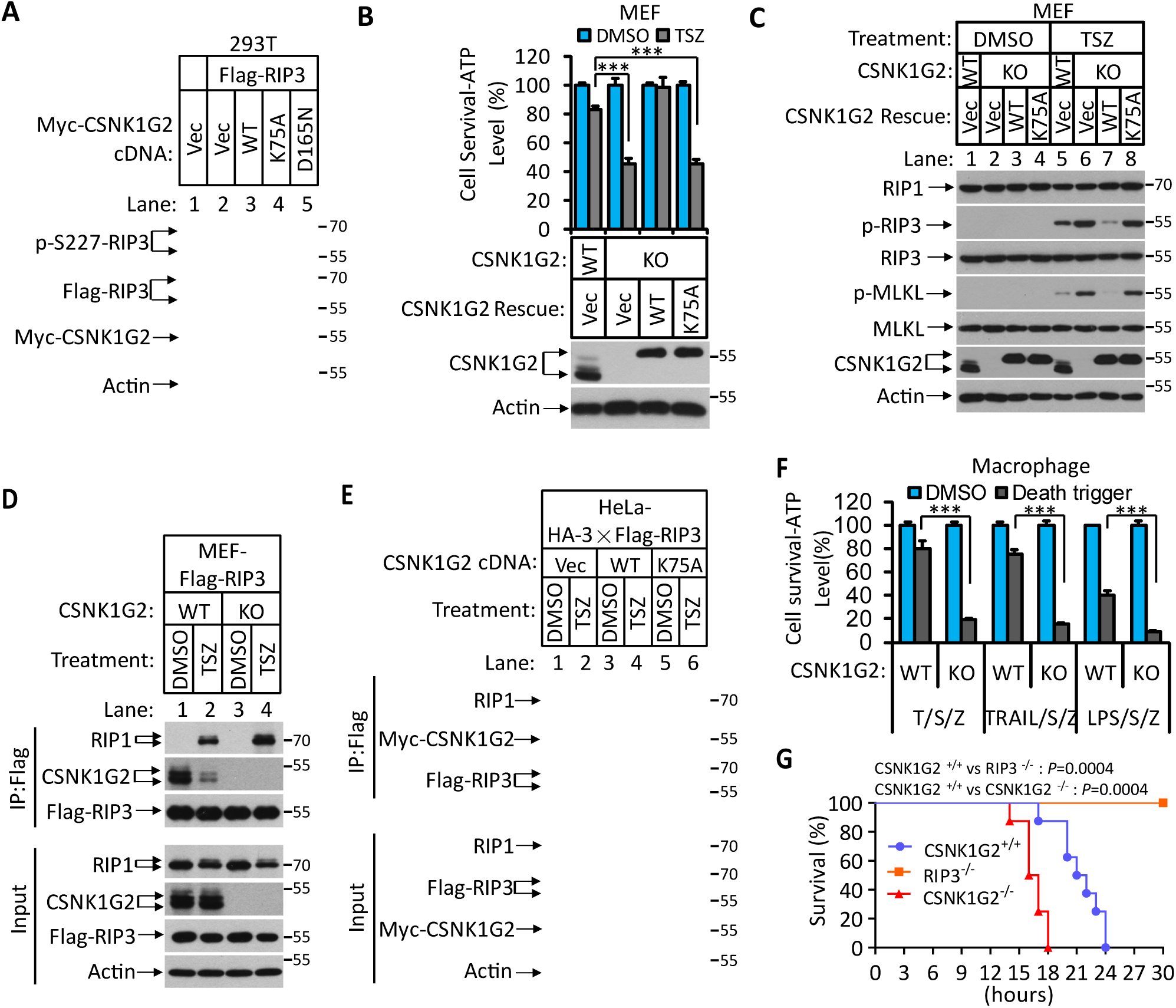
Knockout *CSNK1G2* Accelerates Necroptosis. (A) Western blotting analysis using antibodies against the indicated proteins. Cultured 293T cells were transfected with Flag-tagged RIP3 and the indicated versions of Myc-tagged CSNK1G2, including wild type (WT) and two kinase-dead point mutants K75A and D165N for 20 hrs. Cell extracts were then prepared and used for western blotting analysis. Vec, vector control. Numbers on the right indicate molecular weight markers (kDa). (B) Top: Cell viability as measured by Cell Titer-Glo. Cultured MEF with wild type CSNK1G2 gene (WT) or with their *CSNK1G* gene knocked out (KO) followed by transfection with vector control (Vec) or indicated wild type or a kinase-dead (K75A) mutant CSNK1G2 MEF. The cells were then treated with DMSO or TSZ as indicated for 12 hrs before the intracellular ATP levels were measured by Cell Titer-Glo. T denotes 20 ng ml^-1^ TNF-α; S, denotes 100 nM Smac mimetic; Z denotes 20 μM Z-VAD-FMK. Data are mean ± SD of triplicate wells. \*\*\**P*<0.001. *P* values were determined by two-sided unpaired Student’s *t* tests. Bottom: Aliquots of these treated cells were used for western blotting analysis using an antibody against CSNK1G2. (C) Western blotting of necroptosis activation markers phospho-RIP3 (p-RIP3) and phospho-MLKL (p-MLKL). Cultured MEF cells with indicated CSNK1G2 gene as in (B) were treated with indicated stimuli for 4 hrs before the cell extracts were prepared and subjected to western blotting analysis as indicated. (D and E) Western blotting analysis of RIP3-associated RIP1 and CSNK1G2. Immuno-precipitates using an anti-Flag antibody from extracts of MEF-Flag-RIP3 and MEF (*CSNK1G2^-/-^*)-Flag-RIP3 cells (D) or HeLa-HA-3×Flag-RIP3-Myc-CSNK1G2(WT and K75A) cells (E) treated with the indicated stimuli for 6 hrs were subjected to western blotting analyzing using antibodies as indicated. (F) Cell viability measurement of bone marrow-derived macrophages from the wild type or *CSNK1G2* knockout mice. Macrophages were isolated from the wild type (WT) or *CSNK1G2* knockout mice (KO) and treated with the indicated necroptosis stimuli for 12 hrs, and the cell viability was measured by Cell-Titer Glo. Trail: TNF-related apoptosis-inducing ligand. LPS: Lipopolysaccharide. Data are mean ± SD of triplicate wells. \*\*\**P*<0.001. *P* values were determined by two-sided unpaired Student’s *t* tests. (G) Kaplan-Meier plot of survival of male *CSNK1G2^+/+^*(wild-type), *CSNK1G2^-/-^* (*CSNK1G2* knockout littermates), or *RIPK3^-/-^* (RIP3 gene knockout) mice (n=10 for each genotype, age: 3 months) injected intraperitoneally with one dose of murine TNF-α (300 μg kg^-1^). Body temperature was measured with a lubricated rectal thermometer. Mice with a temperature below 23 °C were euthanized for ethical reasons. See also Figure S1, S2 and S3.

To further explore the mechanism through which CSNK1G2 suppresses necroptosis, MEFs with their endogenous CSNK1G2 knocked out or rescued with wild type or kinase-dead CSNK1G2 cDNA were treated with necroptosis-inducing T/TRAIL/LPS+S+Z, and the necroptosis activation markers phospho-RIP3 (at threonine 231 and serine 232, equivalent to serine 227 of human RIP3) and phospho-MLKL (at serine 345, equivalent to serine 358 of human MLKL) were analyzed by western blotting. Knocking out CSNK1G2 in MEFs resulted in higher levels of phospho-RIP3 and phospho-MLKL (Figure 1C, lanes 5-6 and S2B). Re-introducing wild type, but not kinase-dead mutant CSNK1G2, significantly decreased the levels of phospho-RIP3 and phospho-MLKL when cells were treated with necroptosis inducers (Figure 1C, lanes 7-8).

Necroptosis induced by TSZ is initiated by the formation of necrosome, a protein complex containing both RIP1 and RIP3 (Cho et al., 2009; He et al., 2009; Zhang et al., 2009). MEFs with their CSNK1G2 knocked out showed more RIP1 kinase association with RIP3 (Figure 1D, lanes 2 and 4), indicating that CSNK1G2 suppresses necroptosis by binding to RIP3 and preventing its recruitment to necrosome.

In addition to MEFs, we also investigated the effect of CSNK1G2 in human cells. As shown in Figure 1E, expressing a Myc-tagged human CSNK1G2 reduced RIP1 and RIP3 interaction as measured by a co-immunoprecipitation (co-IP) experiment (Figure 1E, lanes 2 and 4). The kinase-dead mutant of CSNK1G2 (K75A) neither co-IPed with RIP3 nor decreased RIP1-RIP3 interaction (Figure 1E, lanes 5-6, and S2C). Moreover, the wild type CSNK1G2 blocked necroptosis caused by the FKBP-binding small molecule induced dimerization of an FKBP-F36V-RIP3 fusion protein (Li et al., 2020; Orozco et al., 2014), while the kinase-dead CSNK1G2 (K75A) mutants did not, indicating that CSNK1G2 inhibits necroptosis by directly preventing RIP3 activation (Figure S2D).

### *CSNK1G2* Knockout Mice Showed Accelerated TNF-α-Induced Systematic Sepsis

We subsequently knocked out the *CSNK1G2* gene in mice using guide RNA specifically targeted to exon-2 of the *CSNK1G2* gene (Figure S3A). The successful deletion of 56 base pairs of exon-2 in the *CSNK1G2* gene caused the deletion of its N-terminus kinase domain and introduced a new premature stop codon in the remaining mRNA (Figure S3A, S3B and S3D). Knockout of the *CSNK1G2* gene resulted in no detection of CSNK1G2 protein in the testis of these animals (Figure S3C).

Loss of the *CSNK1G2* gene did not affect the development of the knockout mice. However, the bone marrow-derived macrophages (BMDM) from the *CSNK1G2* knockout mice showed enhanced cell death when treated with necroptosis-inducing agents, including T/S/Z, TRAIL/S/Z, or LPS/S/Z (Figure 1F) compared to the BMDM from their wild type littermates. Notably, although the *CSNK1G2* knockout mice looked normal, they died within 18 hours after TNF-α administration, much quicker than their wild type littermates, indicating that TNF-α-induced systematic sepsis was dramatically enhanced (Figure 1G). In contrast, RIP3 knockout mice were totally resistant to such a treatment as previously reported (Newton et al., 2014) (Figure 1G).

### CSNK1G2 Interaction with RIP3 Requires Auto-Phosphorylation of Serine 211 and Threonine 215 Sites

Since the interaction between CSNK1G2 and RIP3 requires the kinase activity of CSNK1G2, we searched for phosphorylation events on these two proteins that might be required for such an interaction. To this end, we immuno-precipitated CSNK1G2-RIP3 complex and by mass spectrometry analysis found three clusters of peptides of CSNK1G2 that contained phosphorylated amino acid residues. These residues were: serine 26 and serine27, serine 211 and thereonine215, and serine 381 (Figure 2A and S4A). We subsequently introduced phosphorylation resistant mutations in these residues and assessed their effect on RIP3 activity. We found that only the S211A/T215A mutant lost the ability to block RIP3 S227 phosphorylation, whereas the other two mutants containing serine 26/serine 27 and serine 381 to alanine mutations still blocked RIP3 S227 phosphorylation as efficiently as the wild type (Figure 2B and S4B). Consistently, re-introducing S211A/T215A mutant CSNK1G2 to the CSNK1G2 knockout MEFs did not restored the necroptosis-inhibiting activity of CSNK1G2 (Figure 2C). Furthermore, compared to wild type CSNK1G2 that efficiently bound to RIP3 and blocked RIP1-RIP3 interaction induced by necroptosis inducer TSZ (Figure 2D, lanes 1-4), S211A/T215A mutant CSNK1G2 lost its ability to bind RIP3 and did not block RIP1-RIP3 interaction in response to TSZ (Figure 2D, lanes 5-6). Moreover, single S211A or T215A mutant showed decreased ability to block RIP3 kinase activity while the double mutant lost all the inhibitory activity, similar to the K75A kinase-dead mutant (Figure S4C). These results suggested that the auto-phosphorylation of serine 211 and threonine 215 contributed to the ability of CSNK1G2 to bind and inhibit RIP3 kinase activity. Not surprisingly, serine 211 and threonine 215 are within the conserved region of CSNK1G2 with amino acid residues between 205 to 240 (human origin) 100% conserved between human, chimpanzee, mouse, and bovine (Figure S4D).

**Figure 2.**
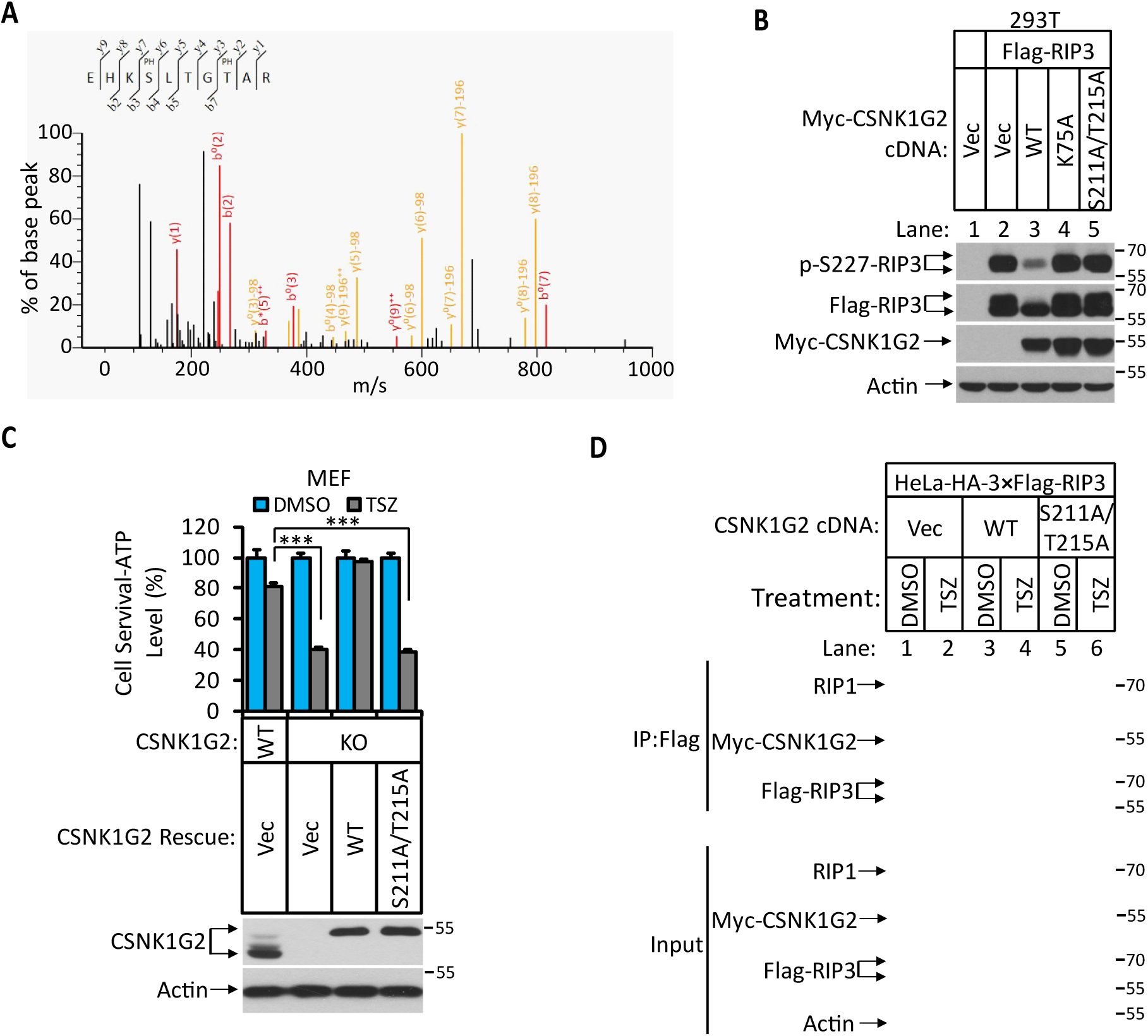
Identification and Characterization of CSNK1G2 Auto-Phosphorylation Sites. (A) MS/MS spectrum of CSNK1G2 phosphorylation sites. The identified phosphorylated peptide to be EHK_P_SLTG_P_TAR with S211 and T215 as the phosphorylated amino acid residues. The b and y type product ions are indicated in the spectrum. Data are related to those in Figure S4A. (B) Effect on RIP3 auto-phosphorylation by co-expression of the indicated version of CSNK1G2. Cultured 293T cells were transfected with Flag-tagged RIP3 cDNA together with indicated Myc-tagged wild type CSNK1G2 (WT), or kinase-dead mutant K75A, or phosphorylation site resistant mutant S211A/T215A for 20 hrs. The cell extracts were then subjected to western blotting analysis using antibodies against phospho-S227-RIP3, Flag-RIP3, and Myc-CSNK1G2 as indicated. (C) Top: Cell viability measurement of effect of phosphorylation sites mutant CSNK1G2 on necroptosis. Cultured wild type MEF (WT), or MEF with their *CSNK1G* gene knocked out (KO) were transfected with either vector control (Vec) or cDNA encoding wild type CSNK1G2 (WT) or phosphorylation sites mutant (S211A/T215A) followed by treatment of DMSO or necroptotic stimuli TSZ as indicated for 12 hrs. The cell viability was measured by Cell-titer Glo. Data are mean ± SD of triplicate wells. \*\*\**P*<0.001. *P* values were determined by two-sided unpaired Student’s *t* tests. Bottom: Aliquots of these treated cells were used to make cell extracts for western blotting analysis using an antibody against CSNK1G2 protein. (D) The effect of CSNK1G2 on RIP1/RIP3 interaction as measured by co-IP. Cultured HeLa-HA-3×Flag-RIP3 cells were transfected with either vector control (Vec) or wild type (WT) or phosphorylation site mutant Myc-CSNK1G2 (S211A/T215A) as indicated. The cells were then treated with DMSO or necroptosis stimuli TSZ for 6 hrs. The cell extracts were prepared and subjected to immunoprecipitation with an anti-Flag antibody. The extracts (Input) and the immuno-precipitates (IP: Flag) were then subjected to western blotting analysis using antibodies as indicated. See also Figure S4.

### Knocking out CSNK1G2 in Testis Cells Significantly Enhanced Their Necroptosis Response

To further study the function of CSNK1G2 *in vivo*, we first measured the expression of this protein in mouse tissues by western blotting. As shown in Figure 3A, CSNK1G2 expression is low in the brain, heart, liver, ovary, and intestine (Figure 3A, lanes 3, 4, 6, 7, 8, 9, 10). There was higher CSNK1G2 presence in lung and spleen (Figure 3A, lanes 2, 5). The highest expression was found in the testis (Figure 3A, lane 1). Immunohistochemical analysis showed that CSNK1G2 was present in the seminiferous tubules of the testis (Figure 3B) and overlapped with that of RIP3, indicating that, like RIP3, it is also expressed in the spermatogonium stem cells and Sertoli cells, two major cell types in the seminiferous tubules (Li et al., 2017). When primary cells from the testis of *CSNK1G2* knockout mice and their wild type littermates were treated with necroptosis stimuli TSZ, significantly more cell death was observed in cells from the *CSNK1G2* knockout testis (Figure 3C). These findings indicate that the function of CSNK1G2 in testis is to block necroptosis in the RIP3-expressing spermatogonium stem cells and Sertoli cells.

**Figure 3.**
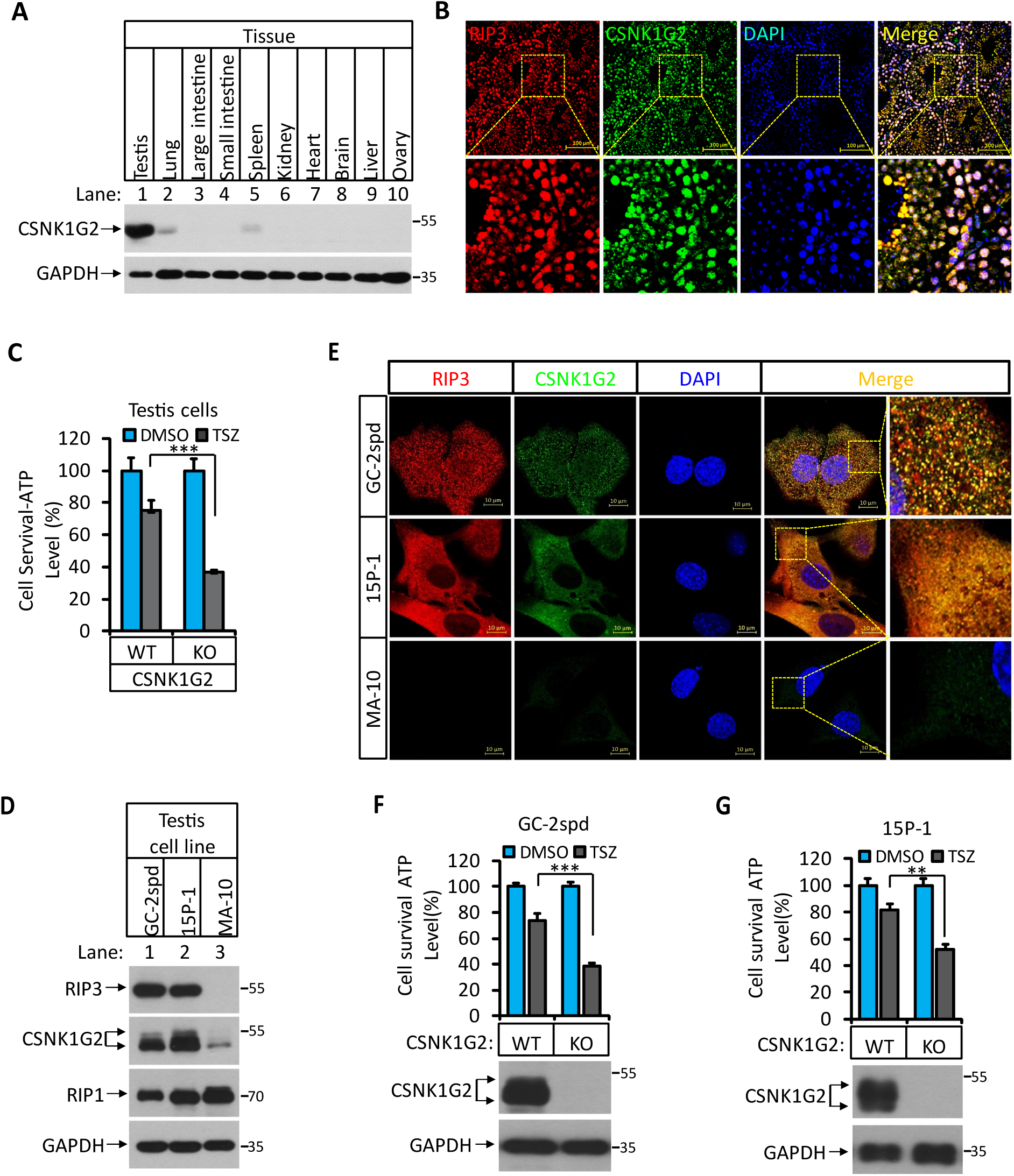
CSNK1G2 and RIP3 are Co-Expressed in the Seminiferous Tubules of the Mouse Testis. (A) The expression of CSNK1G2 in mouse testis, lung, large intestine, small intestine, spleen, kidney, heart, brain, liver and ovary tissues (n=3, 2 months). The indicated tissue extracts were subjected to western blotting analysis using antibodies against CSNK1G2 and GAPDH as indicated. (B) The expression of RIP3 and CSNK1G2 in mouse testis. The testis sections of 2-month old wild type mice (n=3) were stained sequentially with antibodies against RIP3 and CSNK1G2 followed by fluorescent-conjugated secondary antibody. Counterstaining with DAPI, blue. Scale bar on the upper panel is 100 μm. The areas marked by the yellow boxes on the upper panel were shown in the lower panel. (C) The sensitivity of primary cells from the seminiferous tubules of *CSNK1G2*^-/-^ and *CSNK1G2^+/+^* testis to necroptosis induction. The cells from the seminiferous tubules of 2-month old littermates with indicated genotype were isolated and cultured in vitro before treated with DMSO or TSZ as indicated for 12 hrs. The cell viability was then measured by Cell-Titer Glo. Data are mean ± SD of triplicate wells. \*\*\**P*<0.001. *P* values were determined by two-sided unpaired Student’s *t* tests. (D) The expression of RIP3, CSNK1G2 and RIP1 protein in GC-2spd, 15P-1 and MA-10 cells. The extracts from the indicated cultured cells were subjected to western blotting analysis using antibodies against RIP3, CSNK1G2, RIP1 and GAPDH as indicated. (E) Immunofluorescent analysis of RIP3 and CSNK1G2 expression in GC-2spd, 15P-1, and MA-10 cells. The GC-2spd, 15P-1, and MA-10 cells cultured on cover slides were sequentially stained with antibodies against RIP3 and CSNK1G2 followed by secondary antibodies conjugated with red (RIP3) or green (CSNK1G2). Counterstaining with DAPI, blue. Scale bar, 10 μm. (F and G) The effect of CSNK1G2 on necroptosis of GC-2spd and 15P-1 cells. Cultured parental GC-2spd (f) or 15P-1 cells (g) (WT), and GC-2spd or 15P-1 cells with their *CSNK1G2* gene knocked out (*CSNK1G2^-/-^*) were treated with DMSO or TSZ as indicated for 4 hrs. The cell viability was measured by Cell-titer Glo. Data are mean ± SD of triplicate wells. \*\**P*<0.01, \*\*\**P*<0.001. *P* values were determined by two-sided unpaired Student’s *t* tests. Bottom, Immunoblot of CSNK1G2. Cell extracts from aliquots of these cells were also subjected to western blotting analysis using antibodies against CSNK1G2 and GAPDH as indicated, and the results were shown in the bottom.

To further demonstrate necroptosis suppression activity of CSNK1G2 in seminiferous cells in testis, we knocked out the *CSNK1G2* gene in cell lines derived from spermatocyte (GC-2spd(ts)), or Sertoli cells (15p-1) and measured their necroptosis response. Similar to what was seen in testis, GC-2spd and 15p-1 cells expressed both RIP3 and CSNK1G2. In contrast, a cell line from the testosterone-producing Leydig cells did not express RIP3, and CSNK1G2 was present at a much lower level compared to the other two cell lines (Figure 3D and 3E). Similar to what was observed in primary testis cells, GC-2spd and 15p-1 with their *CSNK1G2* gene knocked out showed significantly more death compared to their respective parental cells when treated with necroptotic stimuli (Figure 3F and 3G).

### *CSNK1G2* Knockout Mice Showed Accelerated Male Reproductive System Aging Compared to Their Wild Type Littermates

When male mice reach more than one and a half years of age, their body weight increases, their seminal vesicles grow several fold in both size and weight, and their seminiferous tubules empty as spermatogonium and Sertoli cells undergo necroptosis (Li et al., 2017). We thus measured these aforementioned physiological features of *CSNK1G2* knockout mice and their wild type littermates up to 12 months of age. When these mice were at 2-months of age, their body weight, seminal vesicle size, testis, and appearance of seminiferous tubules were indistinguishable (Figure 4A-4F). However, at 12 months of age, the body weight of *CSNK1G2* knockout mice became significantly higher than their wild type littermates, reaching up to 45 grams on average compared to ∼37 grams for the wild type (Figure 4A). Their seminal vesicles weighted approximately 1 gram, a 10-fold increase from when these mice were 2-months old, and about 2-fold higher than their 12-month old wild type littermates (Figure 4B and 4C). The average size and weight of the testis of 12-month old *CSNK1G2* knockout mice were also significantly smaller than their wild type littermates, and many of their seminiferous tubules were already empty (Figure 4D-4F). At 12 months of age, ∼31% of *CSNK1G2* knockout seminiferous tubules were empty compared to ∼2% empty seminiferous tubules of wild type littermates (Figure 4F).

**Figure 4.**
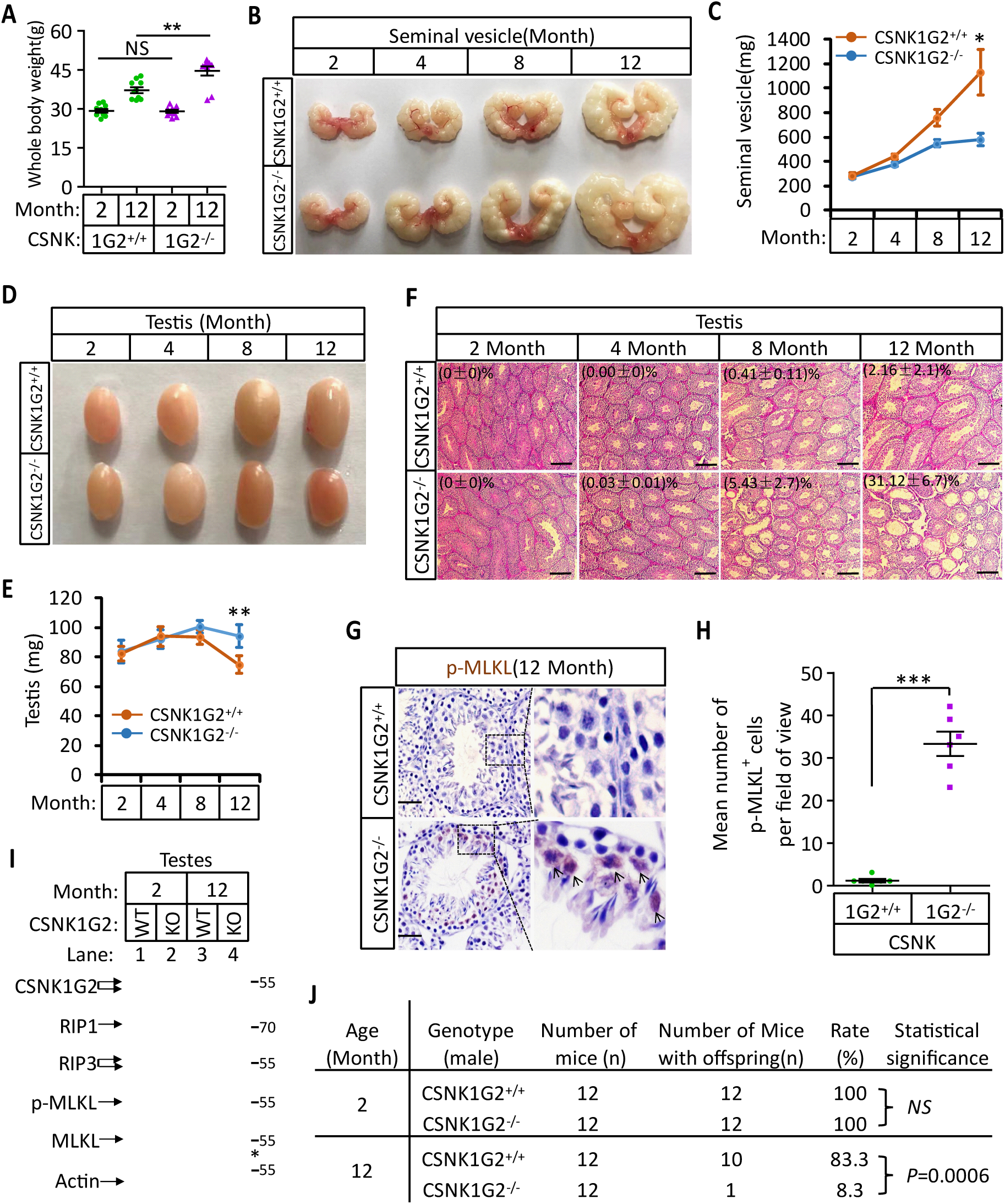
Accelerated Reproductive System Aging in *CSNK1G2^-/-^* Male Mice. (A) Body weights of *CSNK1G2^+/+^* and *CSNK1G2^-/-^* male littermate mice when they were 2- and 12-months old (n=10 for each genotype). (B and C) Macroscopic features (B) and weights (C) of seminal vesicles from *CSNK1G2^+/+^* and *CSNK1G2^-/-^* male littermate mice (n=10 for each genotype) at the indicated ages. (D and E) Macroscopic features (D) and weights (E) of testes from *CSNK1G2^+/+^* and *CSNK1G2^-/-^* male littermate mice (n=12 for each genotype) at the indicated ages. (F) H&E staining sections of testis from *CSNK1G2^+/+^* and *CSNK1G2^-/-^* male littermate mice (n=10 for each genotype) at the indicated ages. The number of empty seminiferous tubules was counted based on H&E staining, and the percentage of empty seminiferous tubules of each group is labeled in the upper left corner of the images. Scale bar, 200 μm. (G and H) Immunohistochemical staining (IHC) of testes from *CSNK1G2^+/+^* and *CSNK1G2^-/-^* male littermate mice (n=6 for each genotype) with phospho-MLKL (p-MLKL) antibody in (G). p-MLKL positive cells were counted in five fields per testis and quantified in (H). Scale bar, 100 μm. (I) Western blotting analysis of extracts from phosphate-buffered saline(PBS) perfused testes of *CSNK1G2^+/+^* (WT) and *CSNK1G2^-/-^* (KO) male littermate mice of 2- and 12-month of age using antibodies against CSNK1G2, RIP1, RIP3, MLKL and phospho-MLKL (p-MLKL) and β-actin as indicated. The number on the right is markers of molecular weight (kDa). Each group was from a pool of three mice. (J) Summary of fertility rates of *CSNK1G2^+/+^* and *CSNK1G2^-/-^* male littermate mice (n=12). Each male mice of 2- or 12-month of age was housed in the same cage with a pairs of 10-week-old wild-type female mice for 3 months; females were replaced every 2 weeks. The number of male mice with reproduction capacity was counted. *P* values were determined using Fisher’s exact tests (unpaired, two-tailed). All quantified data in the figure except (J) represent the mean ± s.e.m. \**P*<0.05, \*\**P*<0.01, \*\*\**P*<0.001. *P* values were determined by two-sided unpaired Student’s *t* tests. NS, not significant.

In addition to the differences in appearance, 12-month-old *CSNK1G2* knockout testis had much more necroptosis activation marker phospho-Serine345-MLKL in their seminiferous tubules. As shown in Figure 4G and 4H, there was prominent staining of phospho-Serine345-MLKL signal in the seminiferous tubules of *CSNK1G2* knockout testis, whereas much less signal was seen in that of wild type littermates. Consistently, the phospho-serine345-MLKL signal was only detected by western blotting in the testis extracts from 12-month old *CSNK1G2* knockout mice (Figure 4I, lane 4). No signal was observed in extracts of young mice (2-month), nor from 12-month old wild type littermates, although the protein levels of RIP1, RIP3, and MLKL were the same (Figure 4I, lanes 1-3).

Finally, we tested the reproductive activity of *CSNK1G2* knockout and their wild type littermates when they were 2 and 12-months of age. As shown in Figure 4J, all twelve 2-month old male mice, regardless of genotype, impregnated their female partners. In contrast, only one of the twelve 12-month old *CSNK1G2* knockout mice was able to produce pubs, whereas ten of the twelve wild type littermates produced progenies.

### Blocking Necroptosis by a RIP1 Inhibitor or *RIP3* Gene Knockout Prevented CSNK1G2-Accelerated Male Reproductive System Aging

To test if the accelerated male reproductive system aging in the *CSNK1G2* knockout mice was indeed due to enhanced necroptosis in the testis of these animals, we fed the *CSNK1G2* knockout animals either a RIP1 kinase inhibitor RIPA-56-containing diet or crossed them with *RIP3* knockout mice to generate *CSNK1G2/RIP3* double knockout mice. As shown in Figure 5A-5C, S5A and S5B, the signs of aging, including body weight gain, an increase of size and weight of seminal vesicles, and a decrease in the size of the testis, were all mitigated when the *CSNK1G2* knockout animals were fed with RIPA-56 or their *RIP3* gene was knockout out. All three physiological features remained at the same level as when the animals were young (3-month old) (Figure S5C).

**Figure 5.**
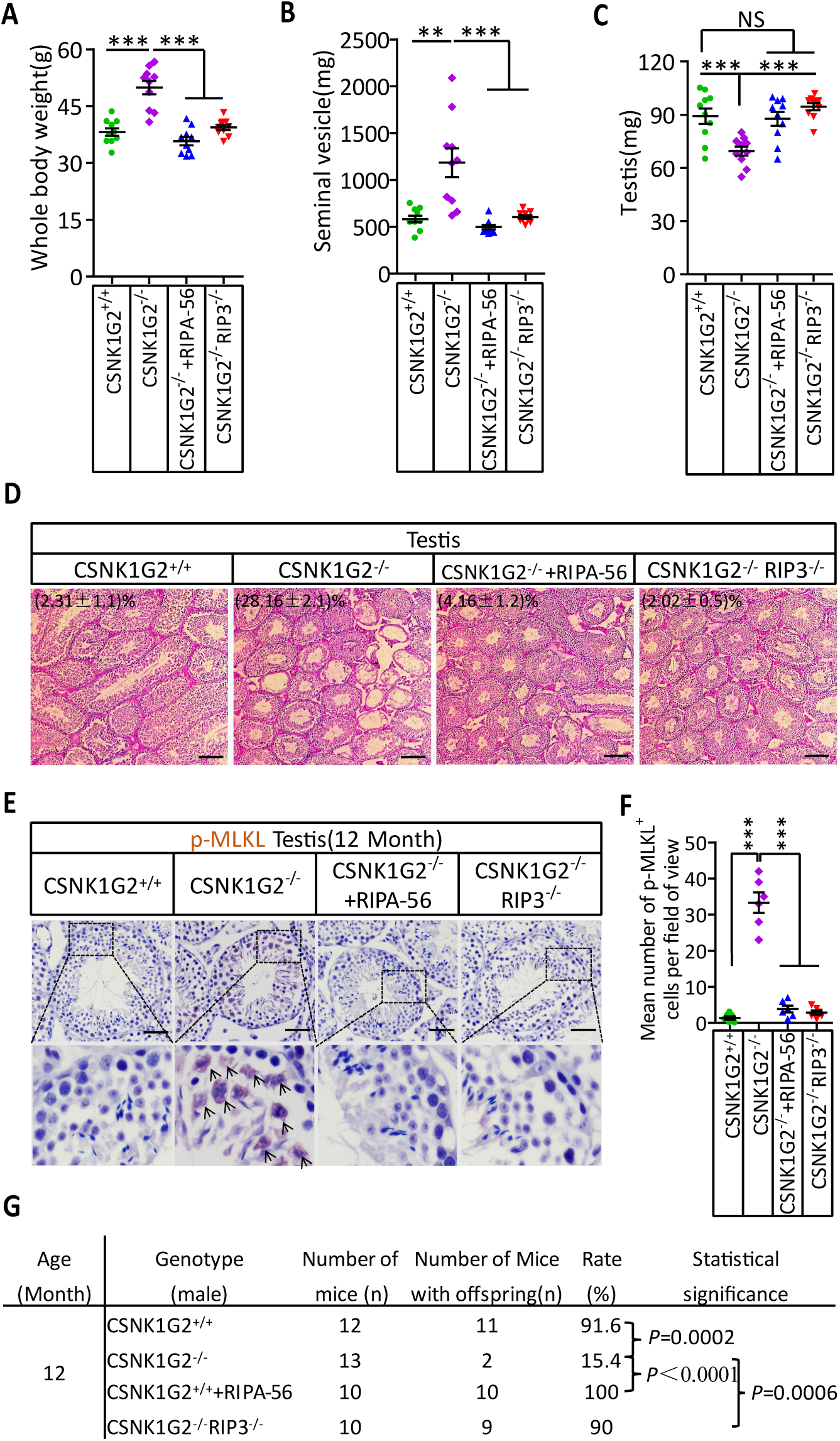
Rescuing the Accelerated Male Reproductive System Aging of CSNK1G2 Knockout Mice with a RIP1 Kinase Inhibitor or RIP3 Knockout. (A) The body weights of 12-month old *CSNK1G2^+/+^*, *CSNK1G2^-/-^*, *CSNK1G2^-/-^* fed with a RIP1 kinase inhibitor (RIPA-56)-containing diet, and *CSNK1G2^-/-^RIP3*^-/-^ male littermate mice (n=10 for each genotype). (B) The weights of seminal vesicles from 12-month old *CSNK1G2^+/+^*, *CSNK1G2^-/-^*, *CSNK1G2^-/-^*+RIPA-56, and *CSNK1G2^-/-^RIP3*^-/-^ male littermate mice (n=10 for each genotype). (C) The weights of testes from 12-month old *CSNK1G2^+/+^*, *CSNK1G2^-/-^*, *CSNK1G2^-/-^*+RIPA-56, and *CSNK1G2^-/-^RIP3*^-/-^ male littermate mice (n=10 for each genotype). (D) H&E staining of testis sections from 12-month old *CSNK1G2^+/+^*, *CSNK1G2^-/-^*, *CSNK1G2^-/-^*+RIPA-56, and *CSNK1G2^-/-^RIP3*^-/-^ male littermate mice (n=10 for each genotype). The number of empty seminiferous tubules was counted based on H&E staining, and the percentage of empty seminiferous tubules was labeled in the upper left corner of the images: scale bar, 200 μm. (E and F) IHC staining of testes from 12-month old *CSNK1G2^+/+^*, *CSNK1G2^-/-^*, *CSNK1G2^-/-^*+RIPA-56 and *CSNK1G2^-/-^RIP3*^-/-^ male littermate mice (n=8 for each genotype) with an anti-phospho-MLKL (p-MLKL) antibody (E). p-MLKL positive cells were counted in five fields per testis and quantified in (F). Scale bar, 100 μm. (G) Summary of the fertility rates of 12-month old *CSNK1G2^+/+^*, *CSNK1G2^-/-^*, *CSNK1G2^-/-^*+RIPA-56, and *CSNK1G2^-/-^RIP3*^-/-^ male littermate mice (n=10 for each genotype). Each male mouse was caged with a pair of 10-week-old wild-type female mice for 3 months; females were replaced every 2 weeks. The number of male mice with reproduction capacity was counted. *P* values were determined using Fisher’s exact tests (unpaired, two-tailed). *CSNK1G2^-/-^*+RIPA-56 mice: *CSNK1G2^-/-^* male mice were fed with AIN93G or AING3G containing RIPA-56 (RIPA-56: 300 mg kg^-1^) for 10 months started when they were 2-month old in an SPF facility. All quantified data in the figure except (G) represent the mean ± s.e.m. \*\**P*<0.01, \*\*\**P*<0.001. *P* values were determined by two-sided unpaired Student’s *t* tests. NS, not significant. See also Figure S5 and S6.

In addition to these gross physical properties, the number of emptied seminiferous tubules in 12-month old mice significantly dropped from almost 30% in *CSNK1G2* knockout mice fed with normal chow diet to about 4% in *CSNK1G2* knockout mice fed with RIPA-56-containing diet (Figure 5D). The number decreased to about 2%, similar to that of 2-month old mice when the mice had their RIP3 gene knocked out (Figure 5D, right panel). The necroptosis activation maker phospho-Serine345-MLKL in the seminiferous tubules of *CSNK1G2* knockout testis was also dropped to almost non-detectable levels in the mice fed with RIPA-56 or had their *RIP3* gene knocked out, in contrast to the prominent signal from 12-month old *CSNK1G2* knockout mice on normal chow (Figure 5E and 5F).

When 12-month old *CSNK1G2* knockout male mice were tested for their reproductive ability, the loss of reproductive function of these mice was almost completely restored when fed with RIPA-56-containing food or had their *RIP3* gene knocked out. All ten 12-month old male *CSNK1G2* knockout mice fed with RIPA-56 containing food produced progenies, and nine of the ten *CSNK1G2*/RIP3 double knockout mice generated pubs when paired with young (2-month old) female partners, whereas only two of the thirteen *CSNK1G2* knockout littermates on chow diet still produced progenies (Figure 5G).

The accelerated aging of the mouse male reproductive system observed in 12-month-old *CSNK1G2* knockout mice also manifested in decreased testosterone levels and increased amount of sex hormone-binding globulin (SHBG), pituitary hormone follicle-stimulating hormone (FSH), and luteinizing hormone (LH) (Figure S6E-S6H). The change in hormonal levels was mitigated when 12-month old *CSNK1G2* knockout animals were fed with RIPA-56 or had their *RIP3* gene knocked out, and all four hormones remained at the same level as when the animals were young (3-month old) (Figure S6A-S6H).

### Human Testes Express CSNK1G2 and Showed Necroptosis Activation Marker When Old

To test if the observed CSNK1G2 expression in mouse testis also occurred in human testis, we analyzed immunohistochemical sections of human testis surgically removed from young men who suffered from severe testicular torsion. As shown in Figure 6A, CSNK1G2 was expressed in the seminiferous tubules of testis in young men, and the expression of CSNK1G2 completely overlapped with RIP3. Interestingly, when we examined the testis of both young and older men, we saw that most of the seminiferous tubules in an 82-year-old’s testis were empty, whereas the seminiferous tubules of a 30-year-old young man were full of cells surrounding vacuolar center where the mature sperms were (Figure 6B and 6C). The seminiferous tubules of an 80-year-old testis also had high signal for phospho-Serine358-MLKL, a marker of necroptosis activation (Figure 6D and 6E). No such signal was detected in the seminiferous tubules of a 30-year old testis (Figure 6D and 6E). These findings indicated that necroptosis-promoted testis aging observed in mice is also conserved in men.

**Figure 6.**
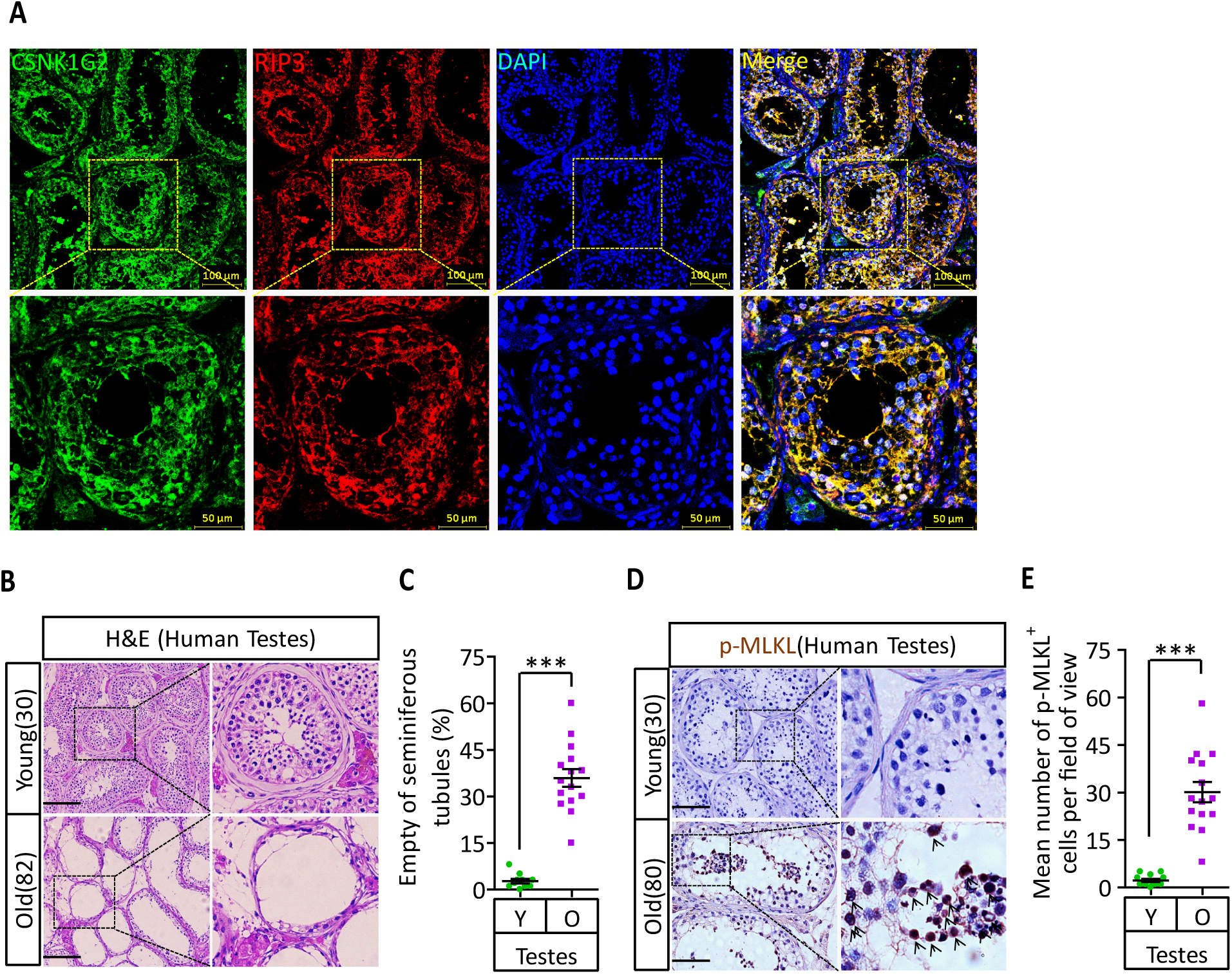
CSNK1G2 Expression and Necroptosis Activation Marker Phosphor-Serine358-MLKL in Human Testes. (A) Expression of CSNK1G2 and RIP3 in human testes. Sections from a testis sample of a 30-years old human patient were stained sequentially with antibodies against CSNK1G2 and RIP3 as indicated followed by green or red fluorescent-conjugated secondary antibodies as indicated. Counterstaining with DAPI, blue. Scale bar, 50/100 μm. Yellow boxes in the upper panels were shown in the lower panels. The experiment was repeated three times with three different patients. (B and C) H&E staining of testes from young and old man. Young man testes (25-30 years, n=10; from testicular torsion necrosis patients) and old man testes (80-89 years, n=15; from prostate cancer patients) were sectioned and stained with H&E in (B). The number of empty seminiferous tubules was counted based on H&E staining and quantification in (C), empty seminiferous tubules were counted in five fields per testis. Scale bar, 100μm. (D and E) IHC of testes from young and old man with phosphor-Serine358-MLKL antibody (p-MLKL). Young man testes (25-30 years, n=10; from testicular torsion necrosis patients) and old man testes (80-89 years, n=15; from prostate cancer patients) were sectioned and stained with an antibody against phosphor-Serine358-MLKL antibody (D). p-MLKL^+^ cells were counted in five fields per testis and quantification in (E). Scale bar, 100μm. All quantified data in the figure represent the mean ± s.e.m. \*\*\**P*<0.001. *P* values were determined by two-sided unpaired Student’s *t* tests.

## DISCUSSION

Necroptosis, although not crucial for normal mammalian development, is known to have critical functions in anti-microbial infection and tissue damage response due to the nature of the inflammation-eliciting necrotic cell death that releases damage pattern recognition (DAMP) signals (Christofferson and Yuan, 2010; Vandenabeele et al., 2010; Wallach et al., 2016). It is thus essential that such a danger signal has to be well-controlled to prevent accidental death. So far, the most dominant negative regulator identified is active caspase-8, which cleaves and inactivates RIP1 and RIP3 kinases, thereby terminating necroptosis (Gunther et al., 2011; Kaiser et al., 2011; Newton et al., 2019; Oberst et al., 2011). Inhibition or genetic inactivation of caspase-8 activates necroptosis both *in vitro* and *in vivo*. Additionally, Ppm1b phosphatase has been proposed to directly dephosphorylate threonine 231 and serine 232 sites of RIP3 to attenuate the recruitment of MLKL (Chen et al., 2015). Moreover, even after MLKL translocated to the plasma membrane, the MLKL-containing membrane fraction lipid rafts can still be removed from the membrane by flotillin-mediated endocytosis or ESCRT-mediated exocytosis before membrane leakage occurs (Fan et al., 2019; Gong et al., 2017; Yoon et al., 2017). All these regulatory events happen after RIP3 kinase is already activated and thus can only serve as “emergency brakes”.

Our current report identified CSNK1G2 as an upstream negative regulator of pre-activated RIP3. CSNK1G2 physically interacts with RIP3 and prevents RIP3 from responding to upstream activating signals, be that TNF-α, TRAIL, or LPS. CSNK1G2’s function appears analogous to cFlip protein for caspase-8 (Chang et al., 2002) or Bcl-2 for Bax/Bak, both of which bind and sequester their pro-death protein targets (Elmore, 2007). The binding of RIP3 by CSNK1G2 requires its auto-phosphorylation on the serine 211 and threonine 215 sites, possibly a structural requirement to form the CSNK1G2/RIP3 complex (Figure S7). Since RIP3 homo-dimerization is a possible requirement for its activation, CSNK1G2, by binding to a monomer RIP3, effectively sequesters RIP3 from activation.

This CSNK1G2-mediated RIP3 suppression is particularly important to attenuate necroptosis from occurring in mouse testis, a cell death program that actively promotes male reproductive system aging (Figure S7). It is thus not surprising that knockout CSNK1G2 resulted in drastically accelerated aging of the testis. CSNK1G2 is also expressed in human testis, and the necroptosis marker phospho-MLKL was present in the old but not young testis of men. This presence is likely the organ-specific aging program of testis mediated by necroptosis of spermatogonium stem cells and Sertoli cells of seminiferous tubules normally suppressed by CSNK1G2, and this phenomenon is evolutionarily conserved between mice and men.

## ACKNOWLEDGMENTS

We thank Mr. Alex Wang for critically reading and editing the manuscript. This work was supported by institutional grants from the Chinese Ministry of Science and Technology and Beijing Municipal Commission of Science and Technology. The funders had no role in study design, data collection and interpretation, or the decision to submit the work for publication.

## AUTHOR CONTRIBUTIONS

D.L. and X.W. designed the research and analyzed the data. D.L., Y.A. and J.G. conducted majority of the experiments. L.L., G.C., and S.C. carried out mass spectrometry analysis. B.D. provided the human samples and helped with data analysis. D.X. and F.W. generated *CSNK1G2^-/-^* mice. D.L. and X.W. wrote the manuscript.

## STAR★METHODS

### LEAD CONTACT AND MATERIALS AVAILABILITY

Further information and requests for reagents should be directed to, and will be fulfilled by the Lead Contact, Xiaodong Wang.

### EXPERIMENTAL MODEL AND SUBJECT DETAILS

#### Mice

The *CSNK1G2* knockout mice were generated using the CRISPR-Cas9 system (Figure S3). The *RIP3^-/-^* (C57BL/6NCrl strain) were kept in our lab(He et al., 2009). *CSNK1G2*^+/-^*RIP3*^+/-^ mice were produced by mating CSNK1G2^-/-^ males with RIP3^-/-^ females. *CSNK1G2*^-/-^*RIP3*^-/-^ mice were produced by mating *CSNK1G2*^+/-^*RIP3*^+/-^ males with *CSNK1G2*^+/-^*RIP3*^+/-^ females. The primers used for genotyping are listed below.

*CSNK1G2*-KO-F: AGGTTTCGCACTCGGATCTCACG;

*CSNK1G2*-KO-R: CCCCGAAGTTCCCACAGCCTATC;

*RIP3*-KO-F: CAGTGGGACTTCGTGTCCG;

*RIP3*-KO-R: CAAGCTGTGTAGGTAGCACATC.

#### Mouse husbandry

Mice were group-housed in a 12 hours light/dark (light between 08:00 and 20:00) in a temperature-controlled room (21.1 ± 1 °C) at the National Institute of Biological Sciences with free access to water. The ages of mice are indicated in the figure, figure legends, or methods. All animal experiments were conducted following the Ministry of Health national guidelines for the housing and care of laboratory animals and were performed in accordance with institutional regulations after review and approval by the Institutional Animal Care and Use Committee at the National Institute of Biological Sciences, Beijing.

#### Human tissues

The research involving human tissue samples were dissected from adult human testicular torsion necrosis and prostate cancer patients (n=10, 25-30 years, testicular torsion necrosis patients; n=25, 80-89 years, prostate cancer patients) were kindly provided from Shanghai Renji Hospital in China and snap-frozen in liquid nitrogen and stored at -80°C. Tissues were cut into appropriately-sized pieces and placed in formalin for preservation. After several days of formalin fixation at room temperature, tissue fragments were transferred to 70% ethanol and stored at 4°C.

The medical ethics committee of the National Institute of Biological Sciences, Beijing, China approved the study.

#### Antibodies and reagents

Antibodies for mouse RIP3 (#2283; WB, 1:1000; IHC, 1:100) were obtained from ProSci. There other antibodies used in this study were anti-GAPDH-HRP (M171-1, MBL, 1:5000), anti-β-Actin-HRP (PM053-7, MBL, 1:5000), anti-Myc-HRP (M192-7, MBL, 1:5000), anti-Flag-HRP (A8592, Sigma-Aldrich), anti-RIP1 (#3493S, Cell Signaling, 1:2000), anti-Human-p-RIP3(ab209384, WB, 1:1000), anti-Mouse-p-RIP3(ab222302, WB, 1:1000), anti-Mouse-MLKL(AO14272B, ABGENT, WB, 1:1000), anti-Mouse-p-MLKL (ab196436; WB, 1:1000; IHC, 1:100), anti-Human-p-MLKL (ab187091; WB, 1:1000; IHC, 1:100), Donkey anti-Mouse, Alexa Fluor 488 (Thermo Fisher, A-21202), Donkey anti-Mouse, Alexa Fluor 555 (Thermo Fisher, A-31570), Donkey anti-Rabbit, Alexa Fluor 488 (Thermo Fisher, A-21206), Donkey anti-Rabbit, Alexa Fluor 555 (Thermo Fisher, A-31572). We collaborated with Epitomics to develop the rabbit monoclonal antibody against CSNK1G2 (ab238121; WB, 1:1000; IHC, 1:100).

### METHOD DETAILS

#### Constructs

psPAX2 and pMD2.G construct were kept in our lab. Full-length human RIP3 and mouse RIP3 cDNA were kept in our lab and subcloned into the pWPI (GFP-tagged) and pCDNA3.1 vector to generate pWPI-Flag-RIP3, pWPI-Flag-mRIP3 and pCDNA3.1-HA-3×Flag-RIP3/RIP3(1-323) construct. Full-length cDNA for human CSNK1A1, CSNK1A1-L, CSNK1D1, CSNK1D2, CSNK1G1, CSNK1G2, CSNK1G3, CSNK1E, CSNK2A1, CSNK2A2 and CSNK2B were RCR-amplified from cDNA library (Sigma) using KOD polymerase (TOYOBO) and subcloned into pcDNA3.1 vector. Using the full-length human and mouse CSNK1G2 cDNA did the truncate pCDNA3.1-Myc-CSNK1G2 (1-408)/CSNK1G2 (192-415)/CSNK1G2 (208-415) and pCDNA3.1-Myc-mCSNK1G2 construct. Using Quickchange Site-Directed Mutagenesis Kit to generate pWPI-Myc-CSNK1G2(K75A)/CSNK1G2 (D165N)/CSNK1G2(S211A)/CSNK1G2(T215A)/CSNK1G2(S211A/T215A) and pWPI-Myc-mCSNK1G2(K75A)/CSNK1G2/CSNK1G2(S211A)/CSNK1G2 (T215A)/CSNK1G2 (S211A/T215A) construct.

The gRNAs for targeting CSNK1G2 (Figure S3) were designed and were cloned into the gRNA-Cas9 expression plasmid pX458-GFP to generate pX458-GFP-mCSNK1G2 construct.

#### Cells

All cells were cultured at 37°C with 5% CO_2_. All cell lines were cultured as follows: HT29 cells were obtained from ATCC and cultured in McCoy’s 5A culture medium(Invitrogen). HEK293T (293T), HeLa, GC-2spd(ts), and 15P-1 cells were obtained from ATCC and cultured in DMEM (Hyclone). NIH3T3-Flag-RIP3(NIH3T3 stably transfected with Flag-RIP3 fused to FKBP-F36V), Mouse embryonic fibroblasts (MEF), MEF (*CSNK1G2^-/-^*), MEF (*RIP3^-/-^*) and HeLa-HA-3×Flag-RIP3(Sun et al., 2012) cells were cultured in DMEM (Hyclone). NIH3T3-Flag-RIP3, HeLa-HA-3×Flag-RIP3 and MEF (*CSNK1G2^-/-^*) cells were infected with virus encoding Myc-CSNK1G2 (WT, K75A and S211A/T215A) or Myc-mCSNK1G2 (WT, K75A, S211A, T215A and S211A/T215A) and GFP-positive live cells were sorted to establish the NIH3T3-Flag-RIP3-Myc-CSNK1G2(WT, K75A), MEF (*CSNK1G2^-/-^*)-Myc-CSNK1G2(WT and K75A) and HeLa-HA-3×Flag-RIP3-Myc-CSNK1G2 (WT and K75A) cell lines. MEF (*RIP3^-/-^*) cells were infected with virus encoding Flag-mRIP3 (WT) and GFP-positive live cells were sorted to establish the MEF (*RIP3^-/-^*)-Flag-RIP3 cell lines. All media were supplemented with 10% FBS (Thermo Fisher) and 100 units/ml penicillin/ streptomycin (Thermo Fisher). MA-10 obtained from ATCC and cultured in DMEM:F12 (Hyclone, additional 20 mM HEPES, horse serum to a final concentration of 15%).

#### Isolation of cells from testes seminiferous tubules

Testes from 8-week old mice were collected using a previously-reported protocol (Chang et al., 2011; Li et al., 2017). Briefly, a testis was placed in Enriched DMEM:F12 (Hyclone) media and placed on ice. After removal of the tunica albuginea of a testis, the seminiferous tubules were dissociated and transferred immediately into 10 mL of protocol enzymatic solution 1. Tubules were incubated for 15-20 min at 35 °C in a shaking water bath at 80 oscillations (osc)/min and were then layered over 40 mL 5% Percoll/95% 1×Hank’s balanced salt solution in a 50 mL conical tube and allowed to settle for 20 min. Leydig cells were removed from the top 35 mL of the total volume of Percoll. The bottom 5 mL of Percoll was transferred to a fresh 50 mL conical tube containing 10 mL enzymatic solution 2. Tubules were incubated for 20 min at 35 °C and 80 osc/min. After incubation, 3 mL charcoal-stripped FBS was immediately added to halt the digestion. The fraction immediately centrifuged at 500×g at 4 °C for 10 min. Pellets were re-suspended in PBS and washed three times, then cultured in DMEM:F12 (15% FBS) medium at 37°C.

#### Cell survival assay

Cell survival assay was performed using Cell Titer-Glo Luminescent Cell Viability Assay kit. A Cell Titer-Glo assay (Promega, G7570) was performed according to the manufacturer’s instructions. Luminescence was recorded with a Tecan GENios Pro plate reader.

#### ELISA

Mice were sacrificed, and blood was clotted for two hours at room temperature before centrifugation at approximately 1,000 × g for 20 minutes. Mice blood sera were collected and assayed immediately or was stored as sample aliquots at -20°C. The testosterone/FSH/LH levels were measured with ELISA kits (BIOMATIK, EKU07605, EKU04284, EKU05693); the SHBG level was measured with an ELISA kit (INSTRUCTION MANUAL, SEA396Mu). The ELISA assays were performed according to the manufacturer’s instructions.

#### CRISPR/Cas9 knockout cells

4 μg of pX458-GFP-mCSNK1G2 plasmid was transfected into 1×10^7^ MEF cells using the Transfection Reagent (FuGENE ®R HD, E2311) by following the manufacturer’s instructions. 3 days after the transfection, GFP-positive live cells were sorted into single clones by using a BD FACSArial cell sorter. The single clones were cultured into 96-well plates for another 10-14 days or longer, depending upon the cell growth rate. The anti-CSNK1G2 immunoblotting was used to screen for the MEF (*CSNK1G2^-/-^*) clones. Genome type of the knockout cells was determined by DNA sequencing.

#### Western blotting

Cell pellet samples were collected and re-suspended in lysis buffer (100 mM Tris-HCl, pH 7.4, 100 mM NaCl, 10% glycerol, 1% Triton X-100, 2 mM EDTA, Roche complete protease inhibitor set, and Sigma phosphatase inhibitor set), incubated on ice for 30 min, and centrifuged at 20,000 × g for 30 min. The supernatants were collected for western blotting. Testis or other tissue were ground and re-suspended in lysis buffer, homogenized for 30 seconds with a Paddle Blender (Prima, PB100), incubated on ice for 30 min, and centrifuged at 20,000 × g for 30 min. The supernatants were collected for western blotting.

#### Immunoprecipitation

The cells were cultured on 15-cm dishes and grown to confluence. Cells at 70% confluence and subjected to indicated treatment for the appropriate time according to different experiments. Then cells were washed once with PBS and harvested by scraping and centrifugation at 800 × g for 5 min. The harvested cells were washed with PBS and lysed for 30 min on ice in the lysis buffer (100 mM Tris-HCl, pH 7.4, 100 mM NaCl, 10% glycerol, 1% Triton X-100, 2 mM EDTA, Roche complete protease inhibitor set, and Sigma phosphatase inhibitor set). Cell lysates were then spun down at 12,000 × g for 20 min. The soluble fraction was collected, and the protein concentration was determined by Bradford assay. Cell extracted was mixed with anti-Flag/Myc affinity gel (Sigma-Aldrich, A2220, A7470) in a ratio of 1 mg of extract per 30 μl of agarose. After overnight rocking at 4 °C, the beads were pelleted at 2,500 × g for 3 min and washed with lysis buffer 3 times. The beads were then eluted with 0.5 mg/mL of the corresponding antigenic peptide for 6 hours or directly boiled in 1× SDS loading buffer (125 mM Tris, pH 6.8, 2% 2-mercaptoethanol, 3% SDS, 10% glycerol and 0.01% bromophenol blue).

#### Harvesting of tissues

Animals were sacrificed and perfused with PBS, followed by 4% paraformaldehyde. Major organs were removed, cut into appropriately-sized pieces, and either flash-frozen in liquid nitrogen and stored at -80 °C or placed in 4% paraformaldehyde for preservation. After several days of 4% paraformaldehyde fixation at room temperature, tissue fragments were transferred to 70% ethanol and stored at 4°C. Blood was collected by cardiac puncture and was allowed to coagulate for the preparation of serum.

#### Immunohistochemistry and immunofluorescence

Paraffin-embedded specimens were sectioned to a 5 μm thickness and were then deparaffinized, rehydrated, and stained with haematoxylin and eosin (H&E) using standard protocols. For the preparation of the immunohistochemistry samples, sections were dewaxed, incubated in boiling citrate buffer solution for 15 min in plastic dishes, and subsequently allowed to cool down to room temperature over 3 hours. Endogenous peroxidase activity was blocked by immersing the slides in Hydrogen peroxide buffer (10%, Sinopharm Chemical Reagent) for 15 min at room temperature and were then washed with PBS. Blocking buffer (1% bovine serum albumin in PBS) was added, and the slides were incubated for 2 hours at room temperature. Primary antibody against p-mMLKL or p-MLKL was incubated overnight at 4°C in PBS. After 3 washes with PBS, slides were incubated with secondary antibody (polymer-horseradish-peroxidase-labeled anti-rabbit, Sigma) in PBS. After a further 3 washes, slides were analyzed using a diaminobutyric acid substrate kit (Thermo Fisher). Slides were counterstained with haematoxylin and mounted in neutral balsam medium (Sinopharm Chemical).

Immunohistochemistry analysis for RIP3 or CSNK1G2 was performed using an antibody against RIP3 and CSNK1G2. Primary antibody against RIP3 was incubated overnight at 4°C in PBS. After 3 washes with PBS, slides were incubated with DyLight-555 conjugated donkey anti-rabbi/mouse secondary antibodies (Life) in PBS for 8 h at 4°C. After a further 3 washes, slides were incubated with CSNK1G2 antibody overnight at 4°C in PBS. After a further 3 washes, slides were incubated with DyLight-488 conjugated donkey anti-mouse/rabbit secondary antibodies (Life) for 2 hours at room temperature in PBS. After a further 3 washes in PBS, the cell nuclei were then counterstained with DAPI (Invitrogen) in PBS. Fluorescence microscopy was performed using a Nikon A1-R confocal microscope.

#### Mating and fertility tests

*CSNK1G^+/+^* and *CSNK1G2^-/-^* male littermate mice were housed in an SPF barrier facility. To score vaginal patency, mice were examined daily from weaning until vaginal opening was observed. The fertility rate of males was determined via a standard method (Cooke and Saunders, 2002; Hofmann et al., 2015; Li et al., 2017) by mating a male with a series of pairs of 10-week-old wild type females for 3 months; females were replaced every 2 weeks (females were either from our colony or purchased from Vital River Laboratory Co(C57BL/6NCrl)). Each litters were assessed from the date of the birth of pups; when pups were born but did not survive, we counted and recorded the number dead pups; for females that did not produce offspring, the number of pups was recorded as ‘0’ (did not produce a litter with a proven breeder male for a period of 2 months). The number of male mice with reproductive capacity was recorded.

#### RIPA-56 feeding experiment (Li et al., 2017)

RIPA-56 in the AIN93G (LAD3001G) at 300 mg/kg was produced based on the Trophic Animal Feed High-tech Co’ protocol. Cohorts of 2-month-old *CSNK1G2^-/-^* male mice were fed with AIN93G or AING3G containing RIPA-56 (RIPA-56: 300 mg/kg) for 10 months in an SPF facility; each male mouse was then mated with four 10-week-old wild-type female mice successively. The number of male mice with reproductive capacity were recorded.

#### Mass spectrometry and data analysis

293T cells were transfected with Myc-tagged CSNK1G2 (WT and K75) for 24 h. Then the cell extracts were prepared and used for immunoprecipitation with an anti-Myc antibody. The immunoprecipitates were washed three times with lysis buffer. The beads were then eluted with 0.5 mg/mL of the corresponding antigenic peptide for 6 hours or directly boiled in 1× SDS loading buffer and subjected to SDS-PAGE. CSNK1G2 bands were excised from SDS-PAGE gel and then dissolved in 2M urea, 50 mM ammonium bicarbonate, pH 8.0, and reduced in 2 mM DTT at 56°C for 30 min followed by alkylation in 10 mM iodoacetamide at dark for 1 h. Then the protein was digested with sequencing grade modified trypsin (Promega) (1: 40 enzyme to total protein) at 37 °C overnight. The tryptic peptides were separated by an analytical capillary column (50 μm × 15 cm) packed with 5 μm spherical C18 reversed phase material (YMC, Kyoyo, Japan). A Waters nanoAcquity UPLC system (Waters, Milford, USA) was used to generate the following HPLC gradient: 0-30% B in 40 min, 30-70% B in 15 min (A = 0.1% formic acid in water, B = 0.1% formic acid in acetonitrile). The eluted peptides were sprayed into a LTQ Orbitrap Velos mass spectrometer (ThermoFisher Scientific, San Jose, CA, USA) equipped with a nano-ESI ion source. The mass spectrometer was operated in data-dependent mode with one MS scan followed by four CID (Collision Induced Dissociation) and four HCD (High-energy Collisional Dissociation) MS/MS scans for each cycle. Database searches were performed on an in-house Mascot server (Matrix Science Ltd, London, UK) against Human CSNK1G2 protein sequence. The search parameters are: 7 ppm mass tolerance for precursor ions; 0.5 Da mass tolerance for product ions; three missed cleavage sites were allowed for trypsin digestion, and the following variable modifications were included: oxidation on methionine, cysteine carbamidomethylation, and serine, threonine, and tyrosine phosphorylation.

### QUANTIFICATION AND STATISTICAL ANALYSIS

All results are representative of three independent experiments. Statistical tests were used for every type of analysis. The data meet the assumptions of the statistical tests described for each figure. Results are expressed as the mean ±s.e.m or S.D. Differences between experimental groups were assessed for significance using a two-tailed unpaired Student’s t-test using GraphPad prism6 and Microsoft Excel 2017. Fertility rate was assessed for significance using Fisher’s exact test (unpaired, two-tailed) using GraphPad prism6 software. The \**P*<0.05, \*\**P*<0.01, and \*\*\**P*<0.001 levels were considered significant. NS, not significant.

---

**Figure S1.**
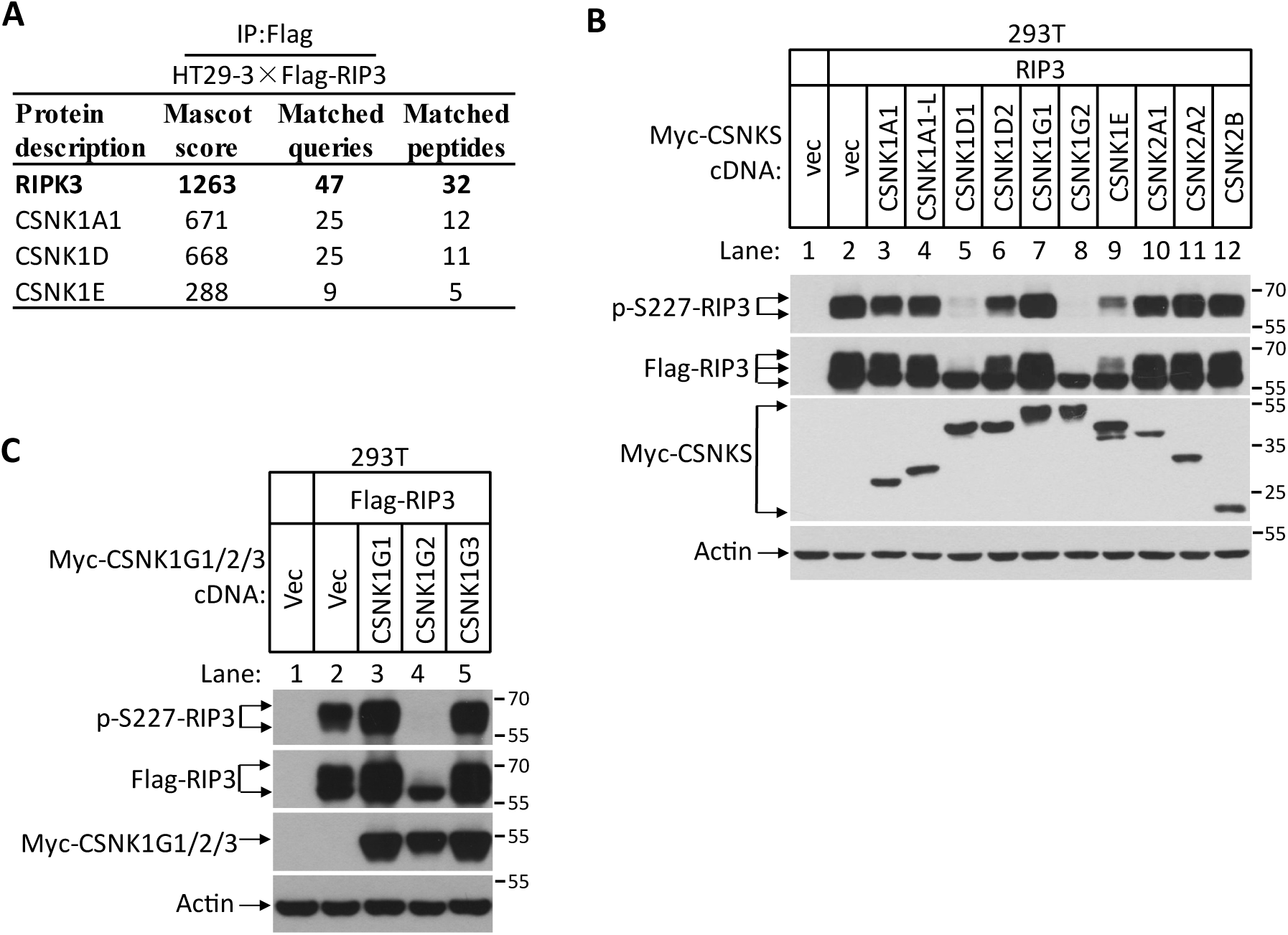
CSNK1G2 Binds and Inhibits RIP3 Kinase Activity to Prevent Necroptosis, Related to Figure 1. (A) CSNK1s associate with RIP3. The HT29-HA-3×Flag-RIP3 cell lysates were immunoprecipitated using anti-Flag resin. The pull-down protein mixture was subjected to mass spectrometry analysis and the identify casein kinase were shown. (B) Measurement of the effect of co-expressed casein kinase members on RIP3 kinase activity. Cultured 293T cells were co-transfected with Flag-tagged RIP3 and indicated Myc-tagged CSNK1A1, CSNK1A1-L, CSNK1D1, CSNK1D2, CSNK1G1, CSNK1G2, CSNK1E, CSNK2A1, CSNK2A2 and CSNK2B for 20 hrs. The cell extracts were then subjected to western blotting analysis using antibodies against Myc-tag, Flag-tag, β-actin, and phosphor-S227-RIP3 as indicated. Numbers on the right indicate molecular weight markers (kDa). (C) Measurement of the effect of co-expression casein kinase 1G members on RIP3 kinase activity. Cultured 293T cells were co-transfected with Flag-tagged RIP3 and indicated Myc-tagged CSNK1G1, CSNK1G2, and CSNK1G3 for 20 hrs. The cell extracts were then subjected to western blotting analysis using antibodies against Myc-tag, Flag-tag, β-actin, and phosphor-S227-RIP3 as indicated.

**Figure S2.**
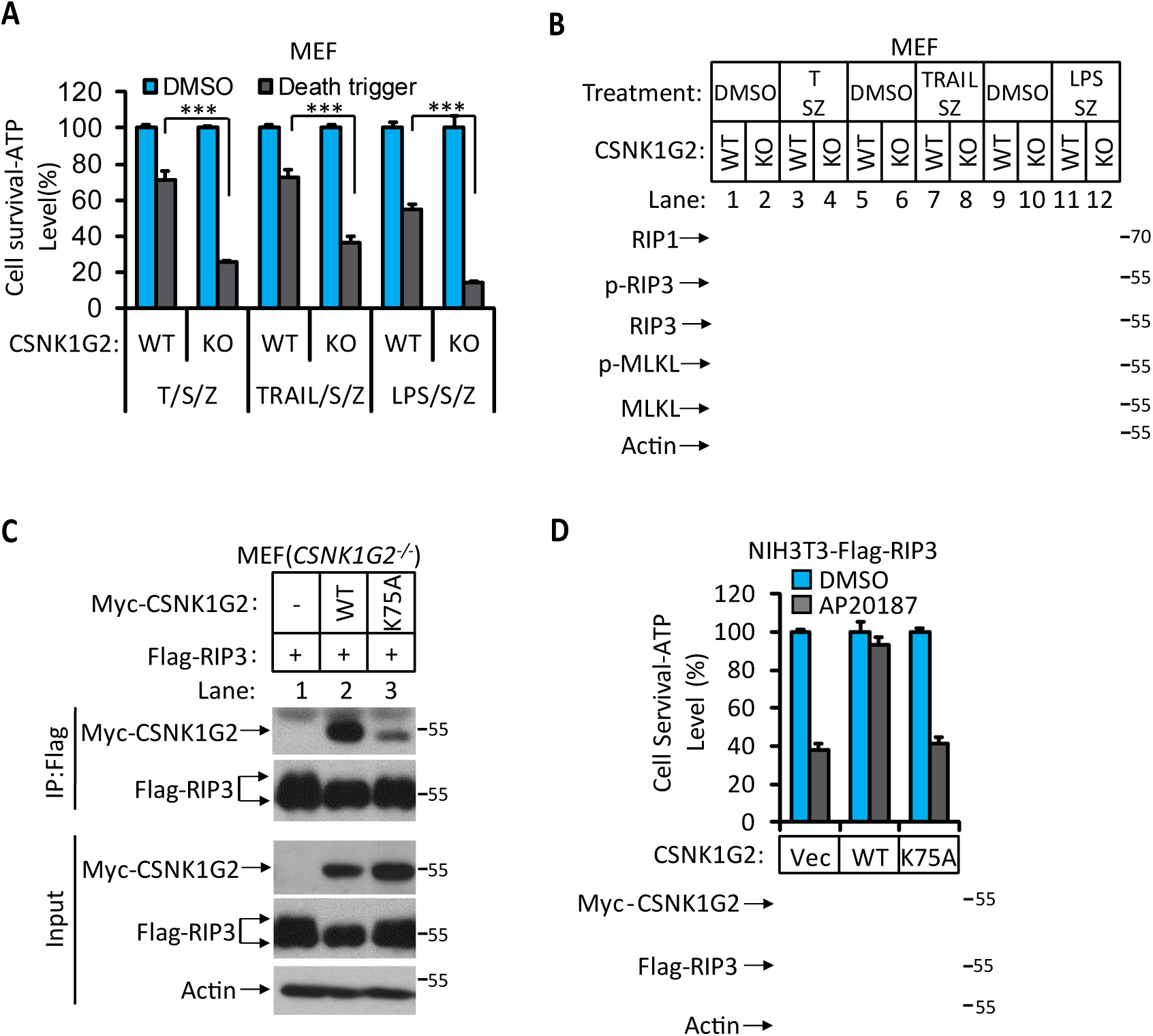
The Kinase Activity of CSNK1G2 is Required for Its Binding to RIP3, Related to Figure 1. (A and B) The necroptotic effect of knockout CSNK1G2 in MEFs. Cultured parental MEF cells (WT) and MEF cells with their CSNK1G2 knocked out (KO) were treated with the indicated necroptotic stimuli for 12 hrs. The cell viability of these necroptotic stimuli-treated cells was then measured using Cell-titer Glo in (A). Data are mean ± SD of triplicate wells. \*\*\**P*<0.001. *P* values were determined by two-sided unpaired Student’s *t* tests. LPS 2 ng ml^-1^, TRAIL 20 ng ml^-1^. The cell extracts were then subjected to western blotting analysis using antibodies against RIP1, phosphor-S227-RIP3 (p-RIP3), β-actin, MLKL, and phosphor-S345-MLKL as indicated in (B). (C) Cultured MEF cells with their *CSNK1G2* gene knocked out (*CSNK1G2^-/-^*) were co-transfected with cDNAs encoding Myc-tagged wild type CSNK1G2 (WT) or kinase-dead mutant (K75A) as indicated and Flag-RIP3 for 24 hrs. The cell extracts were then subjected to western blotting analysis directly (Input), or immunoprecipitation with anti-Flag antibody and the precipitates were subjected to western blotting analysis using antibodies against Myc-tag, Flag-tag, and β-actin as indicated. (D) The effect of wild type and kinase-dead mutant of CSNK1G2 on RIP3 dimerization induced necroptosis. Cultured NIH3T3-Flag-RIP3-Myc-CSNK1G2(Vec, WT, and K75A) cells were treated with the FKBP dimerizer molecule AP20187 for 12h. The cell extracts were subsequently subjected to western blotting analysis using antibodies against Myc-tag, Flag-tag, and β-actin as indicated (lower panel). The cell viability was measured by Cell-titer Glo. Data are mean ± SD of triplicate wells.

**Figure S3.**
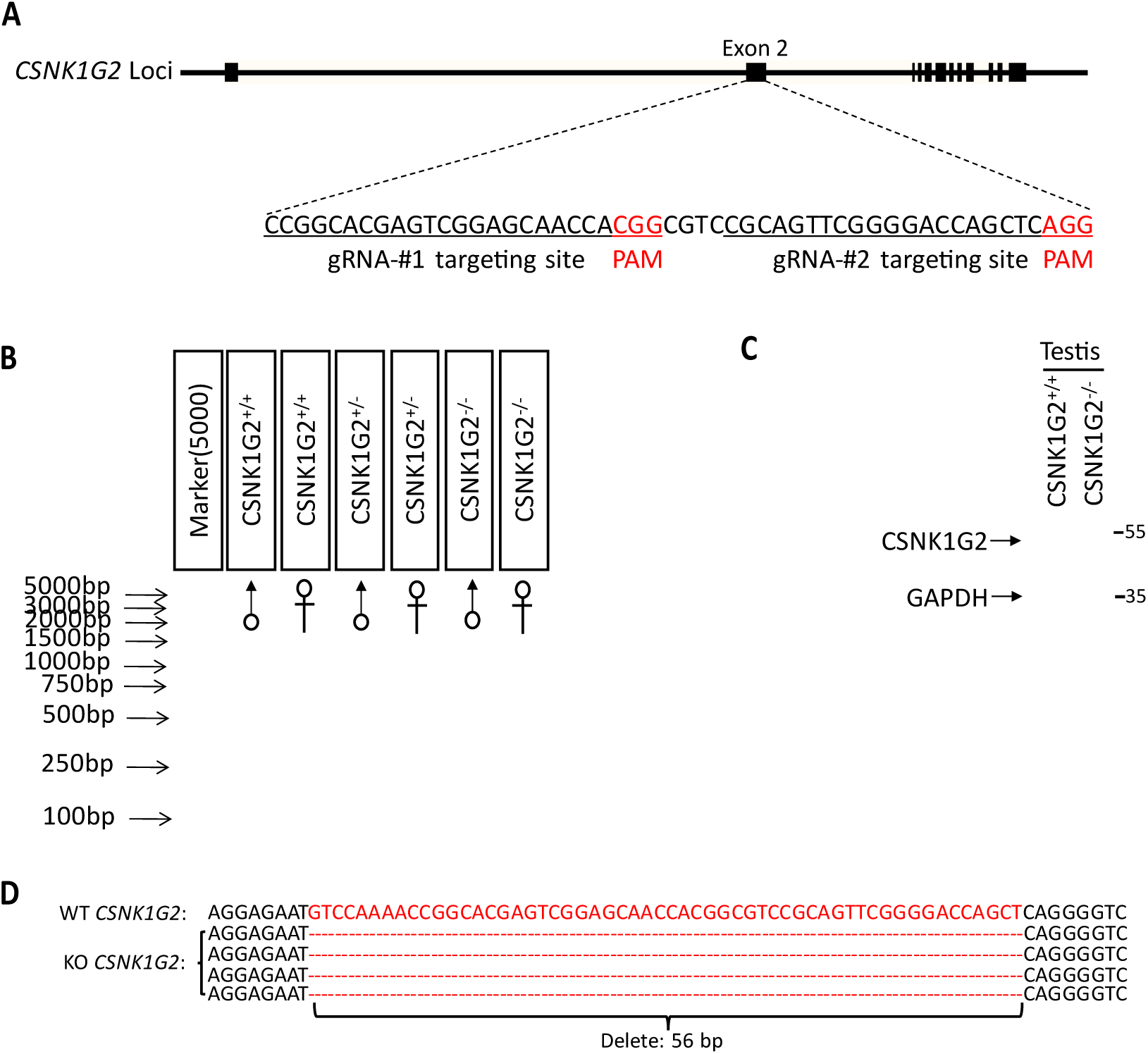
Generation of *CSNK1G2^-/-^* Mice, Related to Figure 1. (A) Schematic of CRISPER-Cas9 strategy for the generation for CSNK1G2 knockout mice. The gene structure of CSNK1G2 and two guide RNA sequences targeting the exon 2 of CSNK1G2 were shown with the PAM sequences highlighted in red. (B) Genotyping results of genomic DNA from *CSNK1G2^+/+^*, *CSNK1G2^+/-^* and *CSNK1G2^-/-^* mice with PCR primers covered the exon 2n targeted region. (C) Immunoblot of CSNK1G2 from testis extracts of 2-minth old *CSNK1G2^+/+^* and *CSNK1G2^-/-^* littermates using antibodies against CSNK1G2 and GAPDH as indicated (n=3). (D) Sequence the genomic of CSNK1G2 knockout mice covering the guide RNA targeted sites. The deleted 56 base pairs of DNA were highlighted in red.

**Figure S4.**
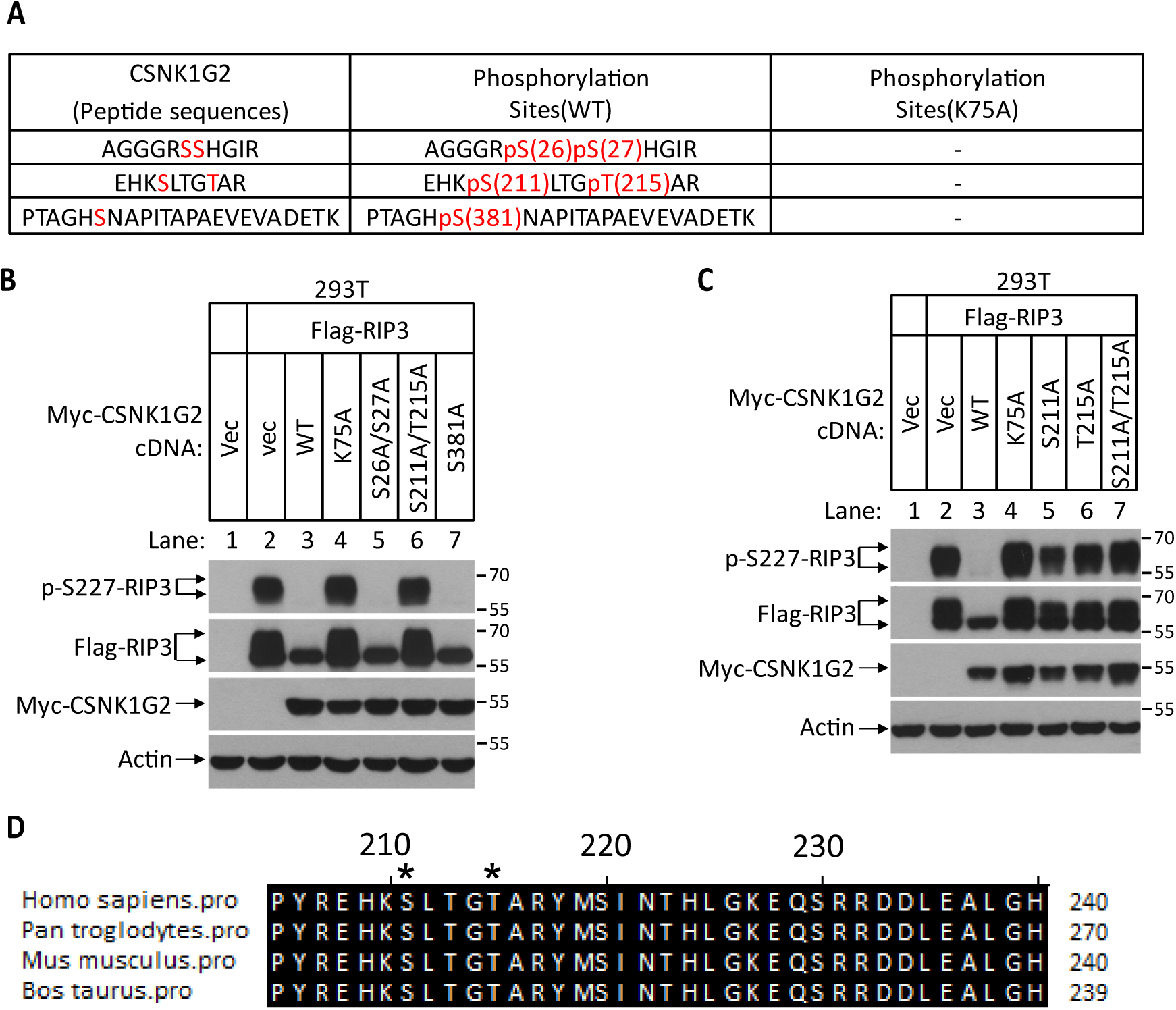
Auto-Phosphorylation Sites on CSNK1G2, Related to Figure 2. (A) Identification of auto-phosphorylation sites on CSNK1G2. Myc-tagged wild type CSNK1G2 or its kinase-dead mutant (WT, K75A) were co-transfected with Flag-RIP3 in 293T cells for 24 hrs. CSNK1G2 was immunoprecipitated using anti-Myc resins. The CSNK1G2 (WT, K75A) bands were excised and analyzed by MS/MS. The identified phosphorylated peptides were shown in the table with the phosphorylated amino acid residues highlighted in red. No phosphorylated peptide was identified in CSNK1G2 (K75A) sample. (B) The effect of phosphorylation sites mutants CSNK1G2 on RIP3 kinase activity. Cultured 293T cells were transfected with vector control (Vec) or Flag-tagged RIP3 and Myc-tagged wild type (WT), or kinase-dead (K75A) or different phosphorylation site mutants as indicated (S26A/S27A, S211A/T215A and S381A) for 20 hrs. The cell extracts were then subjected to western blotting analysis using antibodies against Flag-tag, Myc-tag, phosphor-Serine227-RIP3 and β-actin as indicated. (C) The effect of S211A and T215A mutants of CSNK1G2 on RIP3 kinase activity Cultured 293T cells were transfected with vector control (Vec) or Flag-tagged RIP3 and Myc-tagged wild type (WT), or a kinase-dead mutant (K75A), or S211A or T215A mutants for 20 hrs. The cell extracts were then subjected to western blotting analysis using antibodies against Flag-tag, Myc-tag, phosphor-Serine227-RIP3 and β-actin as indicated. (D) Alignment of amino acid sequences around the auto-phosphorylation sites of CSNK1G2 in four vertebrate species. The Serine 211 and Threonine 215 (human origin) residues are denoted by asterisks (*). The numbers on the right indicate the corresponding histidine in CSNK1G2 of indicated species.

**Figure S5.**
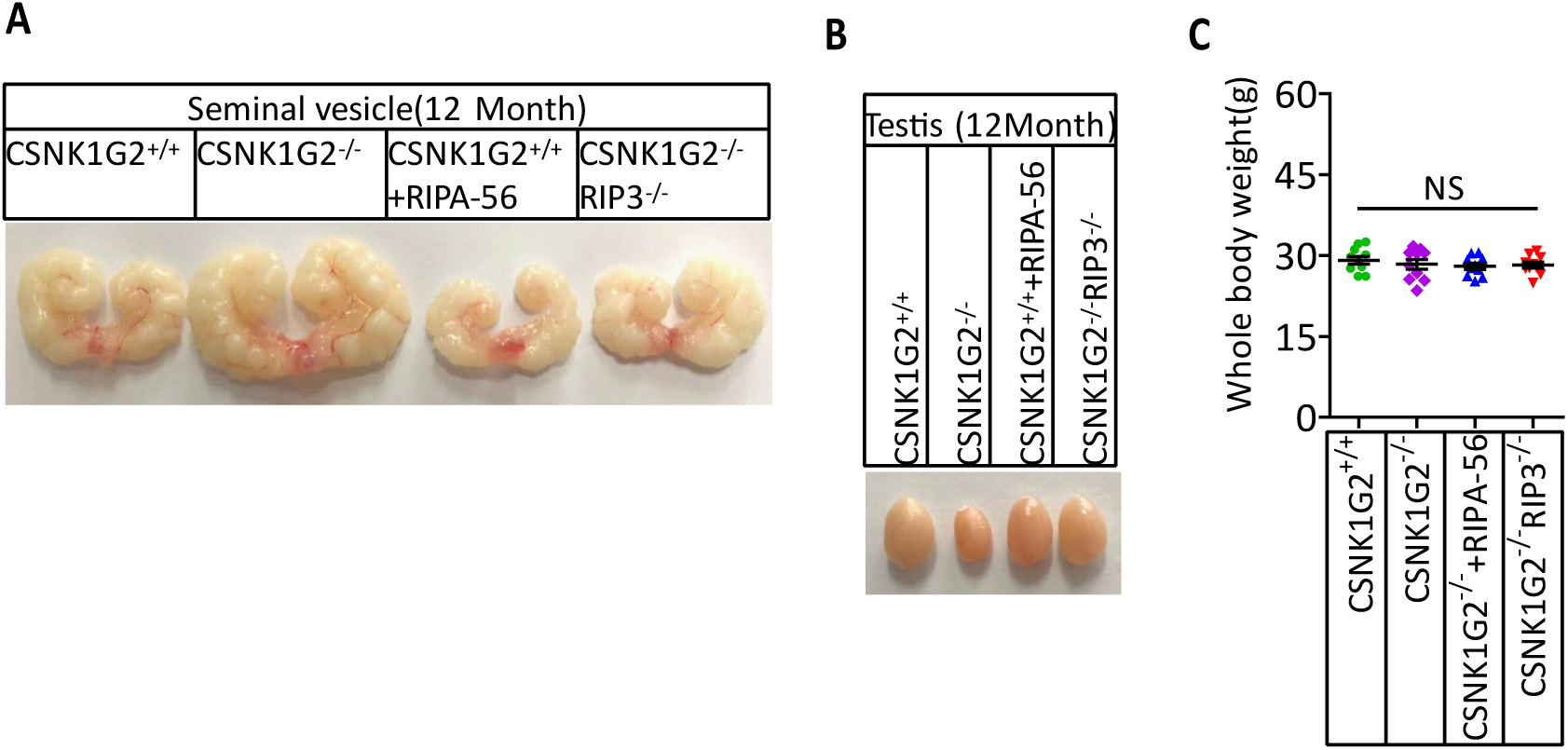
RIP1 Inhibitor-containing Diet or Double Knockout RIP3 Prevents the Appearance of Accelerated Male Reproduction Organ Aging Feature in *CSNK1G2* Knockout Mice, Related to Figure 5. (A) Macroscopic features of a typical 12-month old seminal vesicle from *CSNK1G2^+/+^*, *CSNK1G2^-/-^*, *CSNK1G2^-/-^*+RIPA-56, and *CSNK1G2^-/-^RIP3*^-/-^ male littermate mice (n=10 for each genotype examined). (B) Macroscopic features of a typical 12-month testes from *CSNK1G2^+/+^*, *CSNK1G2^-/-^*, *CSNK1G2^-/-^*+RIPA-56 and *CSNK1G2^-/-^RIP3*^-/-^ male littermate mice (n=10 for each genotype examined). (C) Body weights of 3-month old *CSNK1G2^+/+^*, *CSNK1G2^-/-^*, *CSNK1G2^-/-^*+RIPA-56 and *CSNK1G2^-/-^RIP3*^-/-^ male mice (n=10 for each genotype). *P* values were determined by two-sided unpaired Student’s t-tests. NS, not significant.

**Figure S6.**
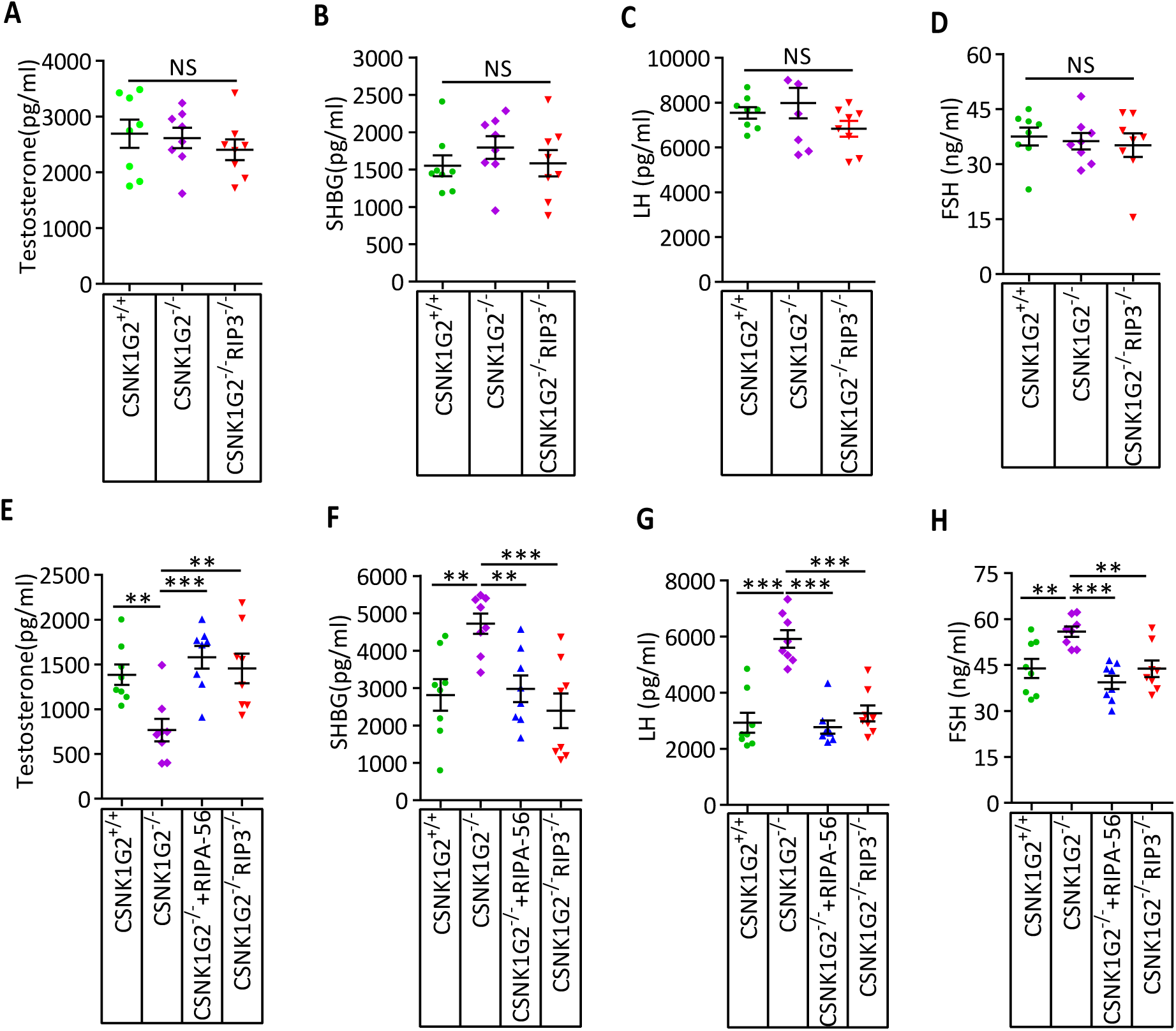
RIP1 Inhibitor-containing Diet or Double Knockout RIP3 Prevents the Hormonal Changes Associated with the Accelerated Male Reproduction System Aging in *CSNK1G2* Knockout Mice, Related to Figure 5. (A-D) Serum hormonal levels of 3-month old *CSNK1G2^+/+^*, *CSNK1G2^-/-^*, *CSNK1G2^-/-^*+RIPA-56 and *CSNK1G2^-/-^RIP3*^-/-^ male littermate mice. Littermates male mice with the indicated genotype were sacrificed, and the indicated hormone levels in serum from were measured using ELISA kit for each hormone (n=8 for each genotype). Data represent the mean ± s.e.m. \*\**P*<0.01, \*\*\**P*<0.001. *P* values were determined by two-sided unpaired Student’s t-tests. *CSNK1G2^-/-^*+RIPA-56 mice: 2-month-old *CSNK1G2^-/-^* male mice were fed with AIN93G or AING3G containing RIPA-56 (RIPA-56: 300 mg kg^-1^) for 1 months in an SPF facility before used. (E-H) Serum hormonal levels of 12-month old *CSNK1G2^+/+^*, *CSNK1G2^-/-^*, *CSNK1G2^-/-^*+RIPA-56 and *CSNK1G2^-/-^RIP3*^-/-^ male littermate mice. Littermates of 12-month old male mice with the indicated genotype were sacrificed, and the indicated hormone levels in serum from were measured using ELISA kit for each hormone (n=8 for each genotype). Data represent the mean ± s.e.m. \*\**P*<0.01, \*\*\**P*<0.001. *P* values were determined by two-sided unpaired Student’s t-tests. *CSNK1G2^-/-^*+RIPA-56 mice: 2-month-old *CSNK1G2^-/-^* male mice were fed with AIN93G or AING3G containing RIPA-56 (RIPA-56: 300 mg kg^-1^) for 10 months in an SPF facility.

**Figure S7. CSNK1G2 Suppresses Necroptosis-Promoted Testis Aging by Binding and Inhibiting RIP3**

Necroptosis induced by TNF-α is initiated by the necrosome formation, a protein complex containing both RIP1 and RIP3. CSNK1G2 blocks necrosome formation through binding RIP3 and preventing RIP3 activation. CSNK1G2 binding to RIP3 is triggered by an autophosphorylation at serine211/threonine 215 sites in its kinase domain. CSNK1G2 knockout mice showed enhanced necroptosis response and pre-maturing aging of their testis.

